# Genetic signatures of human brain structure: A comparison between GWAS and relatedness-based regression

**DOI:** 10.1101/2020.08.07.239103

**Authors:** Bingjiang Lyu, Kamen A. Tsvetanov, Lorraine K. Tyler, Alex Clarke, Cam-CAN, William Amos

**Affiliations:** Centre for Speech, Language and the Brain, Department of Psychology, University of Cambridge, Cambridge, CB2 3EB, UK; Department of Clinical Neurosciences, University of Cambridge, Cambridge, CB2 0QQ, UK; Department of Zoology, University of Cambridge, Downing street, Cambridge, CB2 3EJ, UK

**Author notes:** Correspondence (B.L.), (L.K.T.).

**Keywords:** brain, genetic relatedness, GWAS, linkage disequilibrium

## Abstract

Identifying the genetic variations impacting human brain structure and their further effects on cognitive functions, is important for our understanding of the fundamental bases of cognition. In this study, we take two different approaches to this issue: classical genome-wide association analysis (GWAS) and a relatedness-based regression approach (REL) to search for associations between genotype and brain structural measures of gray matter and white matter. Instead of searching genetic variants by testing the association between a phenotype trait and the genotype of each single-nucleotide polymorphism (SNP) as in GWAS, REL takes advantage of multiple SNPs within a genomic window as a single measure, which potentially find associations wherever the functional SNP is in linkage disequilibrium (LD) with SNPs that have been sampled. We also conducted a simulation analysis to systemically compare GWAS and REL with respect to different levels of LD. Both methods succeed in identifying genetic variations associated with regional and global brain structural measures and tend to give complementary results due to the different aspects of genetic properties used. Simulation results suggest that GWAS outperforms REL when the signal is relatively weak. However, the collective effects due to local LD boost the performance of REL with increasing signal strength, resulting in better performance than GWAS. Our study suggests that the optimal approach may vary across the genome and that pre-testing for LD could allow GWAS to be preferred where LD is high and REL to be used where LD is low, or the local pattern of LD is complex.

## Introduction

The human brain underpins our interaction with the environment through basic sensorimotor processing as well as a wide range of intellectual functions such as attention, memory, decision making and linguistic ability. It is well-known that the structure of the brain is tightly associated with various cognitive functions ^1, 2, 3^, e.g., fluid and crystallized abilities ^4, 5, 6^. However, the genetic variations impacting brain structure, which further affect cognitive functions, are still not fully understood. Resolving the underlying brain-specific genetic signatures will shed light upon our understanding of the fundamental bases of cognition.

Recent genome-wide association studies (GWAS) have revealed genetic variants influencing the volume of both cortical regions ^7^ and subcortical nuclei ^8^. Moreover, attempts have been made to provide a comprehensive description of the genetic determinants of cortical structure which involves measures such as cortical thickness, surface area and volumes ^9, 10, 11^ and white matter microstructure ^12^. GWAS searches genetic variants by testing the association between a phenotype trait (e.g., global or regional brain structure measure) and the genotype of each single-nucleotide polymorphism (SNP) across the genome. Previous studies have revealed that any given brain region’s structural property may be associated with multiple SNPs and that these may be either clustered in one genomic region, or spread across several different chromosomes, suggesting a polygenic architecture ^11^.

In the present study, we adopt a relatedness-based regression approach (REL) ^13^ to search for associations between genotype and brain structure. REL uncovers patterns where individuals with similar genotypes are also similar in their phenotypes and hence uses the same principle as representational similarity analysis, a method widely used in neuroimaging studies ^14^. By summarizing the contribution of multiple SNPs within a genomic window as a single measure, genetic relatedness, statistical power is increased through a reduction in the number of independent tests that are conducted. Furthermore, even when the functional SNP itself has not been sampled, the expectation remains that individuals who are more related are more likely to share a given allele. Consequently, REL has the potential to find associations wherever the functional SNP is in linkage disequilibrium (LD) ^15^ with SNPs that have been sampled. Unlike other methods ^16, 17, 18, 19^ that aggregate multiple SNPs to improve power by reducing multiple-testing burden, REL utilizes the relatedness between individuals, offering a flexible way to control for multiple other variables. For example, it is possible to fit subject age, ethnicity and genome-wide or chromosome-wide relatedness as covariates to control for background shared ancestry and phenotypic changes over a lifetime, while may be challenging in GWAS ^20, 21^.

Here we report the results of a systematic search using both GWAS and REL for genetic variants related to grey matter and white matter phenotypes, as measured by magnetic resonance imaging (MRI). For grey matter, we tested the volume of 55 cortical and subcortical regions (measures of homologous brain areas were averaged across both hemispheres). For white matter, we examined the fractional anisotropy (FA) of 10 white matter tracts which is a measure of integrity (8 within-hemisphere homologous tracts with their FA values averaged across both hemispheres and 2 cross-hemisphere tracts). We also applied principal component analysis (PCA) to all the regional measures of grey matter and white matter separately and searched for the genetic associations of the first two principal components (i.e., PC1, PC2) which could be considered as global measures of brain structure. Finally, we conducted a simulation analysis to evaluate the performance of GWAS and REL with respect to different levels of LD.

## Materials and methods

### Cambridge Centre for Ageing and Neuroscience (Cam-CAN) project

All the data analysed in this study are from the Cam-CAN project (www.cam-can.com) which focuses on understanding the lifespan neurocognitive development. It consists of a population-based sample of 708 healthy participants, age range 18-88 years with a wide range of epidemiological, cognitive, genetic and neuroimaging data (for more details see ^22^). The Cam-CAN project was approved by the Cambridgeshire 2 (now East of England - Cambridge Central) Research Ethics Committee. All participants provided written informed consent.

### Genetic data

Whole-genome SNP data were collected from 708 individuals. DNA was genotyped using Illumina Infinium OmniExpressExome arrays. This chip covers >960,000 SNPs spread through the genome, capturing a large proportion of common variation. The raw genotype data were filtered in GenomeStudio according to standard procedures ^23^. Additional quality control checks were performed in PLINK ^24^ (SNPs were removed if Hardy Weinberg p < 1×10^−6^, missingness > 0.05, or minor allele frequency < 0.01; individuals were removed if total SNP missingness > 0.05, or where multidimensional scaling indicated non-European origin). Lastly, ambiguous SNPs were removed (i.e., A-T or G-C SNPs). After quality control, the dataset included 628,511 directly genotyped SNPs from 634 CamCAN individuals.

### Brain imaging data

Brain imaging data were collected from all 708 participants using a Siemens 3T TIM TRIO (Siemens, Erlangen, Germany) magnetic resonance imaging (MRI) scanner at the Medical Research Council Cognition & Brain Sciences Unit (MRC CBU), Cambridge, UK. Two widely used metrics were adopted to characterise individual brain structure: regional grey matter volume and white matter tract FA. See ^25^ for more details about MRI processing pipeline and extraction of the brain structural measures mentioned below.

Regional grey matter volume characterises the distinct morphological signature of the size of anatomical regions of interest (ROIs) in individuals. Fractional Anisotropy (FA) reflects the integrity of white matter tracts reconstructed in diffusion tensor imaging (DTI) which maps the tractography in the brain ^26^. FA was obtained by quantifying the anisotropy of the movement of water molecules constrained by the fibre bundles as a scalar that ranges from 0 (completely isotropic) to 1 (completely anisotropic).

Regional grey matter volume was estimated using 1mm^3^ isotropic T1- and T2-weighted images which were acquired by a MPRAGE sequence (TR = 2250 ms, TE = 2.98 ms, TI = 900 ms, FA = 9°, FOV = 256 x 240 x 192 mm^3^) and a SPACE sequence (TR = 2800 ms, TE = 408 ms, FA = 9°, FOV = 256 x 256 x 192 mm^3^). The T1- and T2-weighted images were co-registered to the Montreal Neurological Institute (MNI) template and combined to segment the brain into images of different tissues. A group template image was created by conducting diffeomorphic registration (DARTEL) using the grey matter images. We adopted the Harvard Oxford Atlases (HOA) which defines 116 anatomical ROIs that cover all the grey matter in the brain (Fig. 1a). Results of six limbic and subcortical ROIs were not used due to technical issues, thus the regional grey matter volume was only calculated for the remaining 110 ROIs which were further normalized by the total intracranial volume to control for individual differences. Regional volume of homologous areas was averaged across both hemispheres, resulting in 55 phenotype measures (see Table S1 for the full list of all ROIs) to be tested in GWAS and REL.

**Fig. 1.**
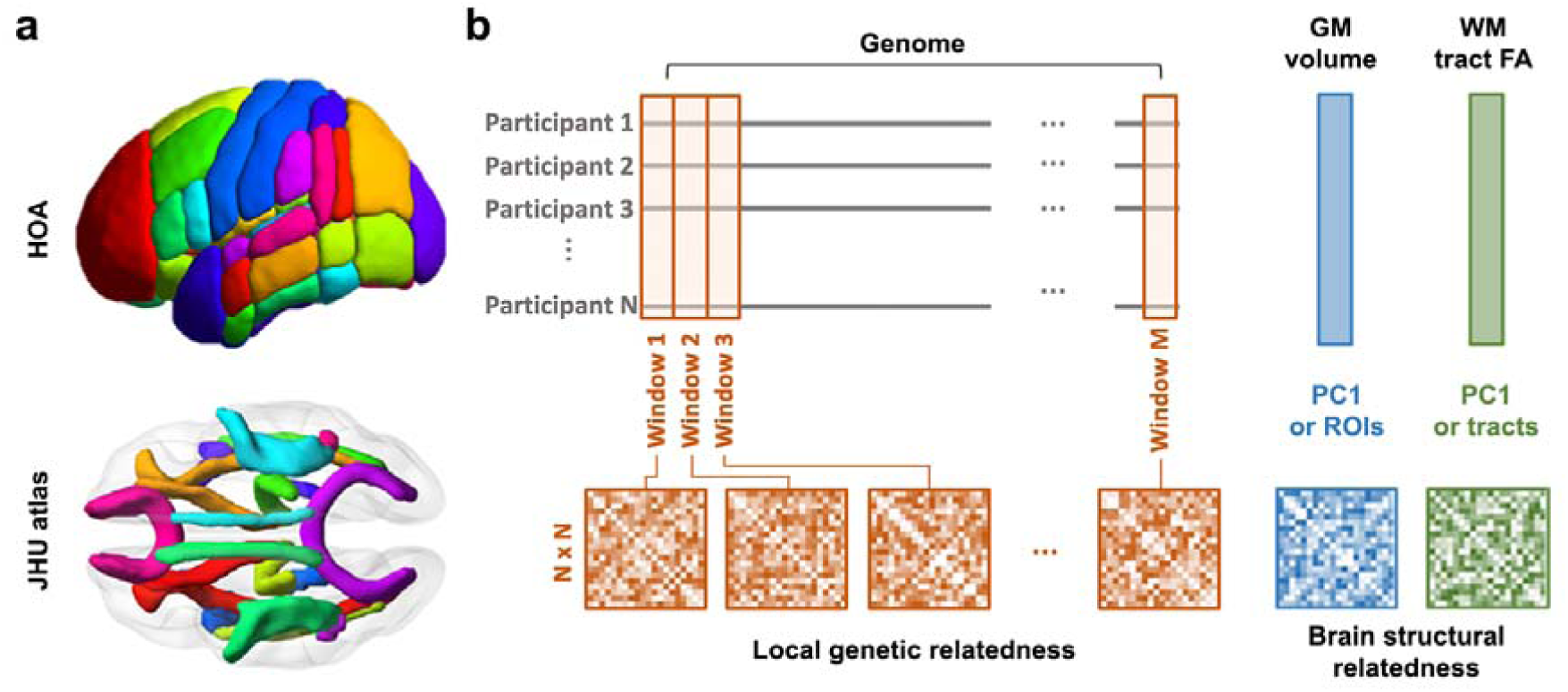
(a) Grey matter ROIs of the Harvard Oxford Atlas (HOA) and white matter tracts defined by the John Hopkins University (JHU) altas. (b) Illusration of the calulation of local genetic relatedness and brain structural relatedness. Elements in the upper triangular part of each relatedness matrix (except for the diagonals) were vectorised and used in the regression model. N: number of participants, GM: grey matter, WM: white matter, FA: fractional anisotropy, PC1: first principle compent.

White matter FA was estimated using 2mm^3^ isotropic diffusion-weighted images acquired with a twice refocused-spin-echo sequence (30 diffusion gradient directions with each for b-values 1,000 and 2,000 s·mm^−2^, three images with a b-value of 0, TE = 104 ms, TR = 9.1 s, 66 axial slices, FOV = 192 × 192 mm^2^). DWI images were corrected for eddy current and head motion, then skull-stripped and co-registered with individual T1-weighted image, finally fitted to a non-linear diffusion tensor model. FA was computed at each voxel using the tensor’s eigenvalues. We adopted the John Hopkins University (JHU) white matter tractography atlas which covers 20 white matter tracts in the brain (Fig. 1a). Data of two white matter tracts were unavailable due to technical issues, thus the structure of white matter was characterised by the mean FA value of each of the 18 available tracts. Apart from the two cross-hemisphere tracts, FA values of the 16 within-hemisphere tracts were averaged across hemispheres (see Table S2 for the full list of all tracts).

Participants who lacked relevant MRI scans or had poor-quality data were excluded. Any participant with measure that was 3 standard deviations away from the group mean was also excluded. Among the 708 participants, 586 of them had data for the calculation of regional grey matter volume and 577 of them had data for the calculation of white matter FA. All MRI data were analysed using the SPM12 (www.fil.ion.ucl.ac.uk/spm) and FSL (fsl.fmrib.ox.ac.uk). To control for the effects of age and gender, we conducted a linear regression for each grey matter or white matter phenotype measure with both age and gender as predictor variables. The standardized residuals were used in further GWAS and REL analyses. PCA was applied to these age and gender corrected brain measures of grey matter and white matter separately. Their first two principle components (PC1 and PC2) were not linear correlated with either age or gender. As global measures, PCs of grey matter and white matter were also adopted to find genetic associations using GWAS and REL.

### GWAS

GWAS was conducted using GEMMA ^27^, based on a total of 628,511 biallelic SNPs. Significance was determined according to the Wald p-value returned in the fitted model. We tested individual grey matter ROIs and whiter matter tracts as well as their first two PCs. Bonferroni correction was applied to correct for the number of SNPs tested for each brain measure, resulting in a genome-wide significance threshold of P = 7.96 x 10^−8^.

### Relatedness-base regression

REL was conducted by fitting the multiple linear regression in the following form:

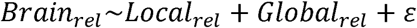

Each pairwise comparison between the brain measure of two individuals yielded a similarity score Brain_rel_ which is the response variable. Local relatedness (Local_rel_) was calculated using SNPs in a genomic window with fixed size, while global relatedness (Global_rel_) was calculated using all the SNPs (outside the genomic window) on the chromosome where the current genomic window is located ^13^. We prefer to use same chromosome markers rather than all other markers because we are not interested in relatedness *per se*, but more to control for background stochastic variation in the probability of markers being identical.

For each grey matter ROI and each white matter tract, we removed linear age- and gender-related changes in phenotype by extracting standardised residuals from a regression with age and gender as the predictor variables. These standardized residuals were then used as input variable in a PCA. The first two components of the white matter tract FA explained 38.8% and 12.8% of the total variance separately, while those of the regional grey matter volume explained 22.8% and 6.2% of the total variance separately. For each brain measure (either PC score or regional measure), we first calculated the absolute difference between individuals, which were then subtracted from the maximal absolute pair-wise difference to obtain the brain relatedness, i.e., Brain_rel_ (Fig. 1). Both local and global relatedness were estimated using the same algorithm ^28^. Non-overlapping genomic windows were constructed with sizes of either 50kb or 100kb (Fig. 1). For 50kb size, there are 52,210 windows with an average of 11.5 ± 7.2 SNPs (mean ± std) per window. For 100kb size, there are 26,383 windows in total with 22.6 ± 12.9 SNPs per window.

Significance of the relationship between the brain measure and the local genetics was determined by a variation of the Mantel test ^13^. Specifically, for each genomic window, we permuted the variable local_rel_ by re-calculating it with a randomised order of individuals in each permutation, which yields an Akaike Information Criterion (AIC) value. P-values were calculated as the ratio of permutations that resulted in a lower AIC compared with the non-permutated value. For the purpose of improving computational efficiency, the permutation tests were terminated when 5 extreme AIC values (i.e., lower than the original value) were obtained or where the total permutation number reached 2 million, thus the lower limit of p-value is 5 x 10^−7^. Note that p-values will tend to be slightly conservative if the permutation test was ended with more than 3 extreme AIC values. Bonferroni correction was applied to correct for the number of windows for each brain measure, resulting in genome-wide thresholds of P = 9.58 x 10^−7^ and P = 1.90 x 10^−6^ for 50kb- and 100kb genomic windows separately.

### Simulation for the comparison between GWAS and REL

We were interested to compare the performance of GWAS and REL on our data. In any given genomic region, performance is likely to be influenced by the LD present, since this will determine the extent to which the genotype at one locus carries information at a nearby, potentially unsampled, second locus. To explore this aspect, we conducted a simulation-based analysis, using the Python package ‘moments’ ^29^ to estimate LD for the covariance of genotypes, D, from unphased data.

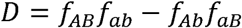

where A/a and B/b are the alleles at two different loci.

To avoid biases due to variation in information content and a possible correlation between SNP density and the variance in estimate of D, for our study we selected 100kb windows containing exactly 25 SNPs. Within each qualifying window we calculated all pair-wise LD values as D^2^ and used the average to represent the LD of the window as a whole. Note that there existed SNP pairs with a negative D value in some genomic windows, which may cancel out the overall within-window effects when both positive and negative D values are averaged, thus we used D^2^ instead of D. Across the genome we found 26,383 windows (i.e., the same 100kb-window division for empirical data) and from these selected 100 that that covered the full observed range of values for mean D^2^.

We next simulated a genotype-phenotype association. To do this, one of the 25 SNPs was selected from a window and designated the ‘functional’ SNP. To make it functional, an artificial phenotype was created by re-assigning the PC1 values for regional grey matter volume across individuals according to the genotype of each individual at the functional SNP. The strength of association could be varied by adding Gaussian noise 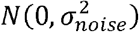 to the artificial phenotype measure, where *σ*_*noise*_ =*σ*_*phenotype*_ /SNR. SNR is the signal-to-noise ratio (SNR) and was set in the range 0.1 to 1 to vary the strength of the signal present (see an example of simulated signal with SNR = 1 in Fig. 2A).

**Fig. 2.**
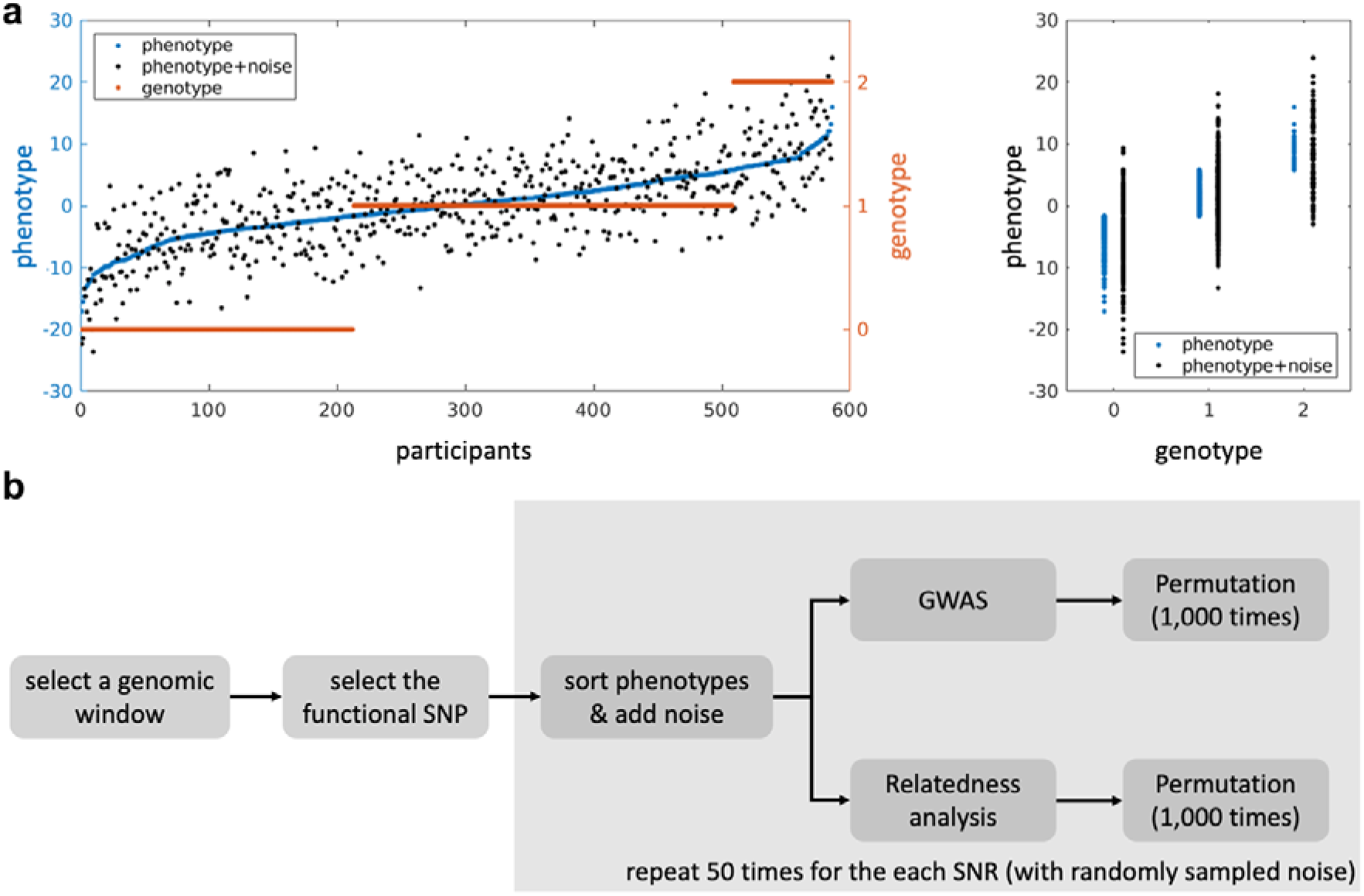
Illustration of simulation for the comparison between GWAS and REL. (a) Left: Individual phenotype measures of GM PC1 (orange dots) were sorted acorrding to the genotypes (blue dots) of the functional SNP in a genomic window. Gaussian noise was then added to obtain the simulated phenotype (black dots) with a certain SNR (i.e., σ_*phenotype*_/ σ_*noise*_, SNR = 1 in this example). Right: comparison between the true signal (sorted phenotypes) and the simulated signal with noise. (b) Schematic diagram pipeline of the simulation for both GWAS and REL.

For each genomic window, simulations was conducted twice with different functional SNPs:

1. with the SNP in the highest overall LD with the other SNPs within the window (measured by the sum of absolute D between the functional SNP and the other SNPs) i.e., Max_LD_ fSNP;
2. with the SNP in the lowest overall LD, i.e., Min_LD_ fSNP. Note that the functional SNP was excluded in the following GWAS and REL. Simulation of one genomic window was repeated 50 times with different Gaussian noise sampled with the same SNR. For both GWAS and REL, significance was determined through permutation test. We ran 1,000 permutations for both GWAS and REL with a randomised order of individuals in each permutation (see Fig. 2b for a schematic diagram of the simulation pipeline). For GWAS, genotype of each SNP was permutated. For REL, both local and global relatedness was recalculated after shuffling the individuals in the cohort. P-value is calculated as the ratio of permutations that return a parameter estimate larger than the original value.

## Results

Below we present the results of two tests for genotype-phenotype associations, the pairwise relatedness analysis, REL, and a classical genome-wide association analysis, GWAS. Both approaches were applied both to each individual measure (grey matter, n = 55 ROIs; white matter, n = 10 tracts) and also to the four metavariables PC1_GREY_, PC2_GREY_, PC1_WHITE_ and PC2_WHITE_.

### GWAS

Applying conventional GWAS, we identified ten genome-wide significant loci associated with either individual measures of grey matter volume or individual white matter tract FA. However, no hit was identified for either PC1 or PC2 for either of these two types of brain structure measures. Fig. 3 summarises the hits for individual measures (see Table S3 for the full description of all the significant loci). For the regional grey matter volume, rs872376 on chromosome 1 is associated with parietal operculum cortex, rs36016914 on chromosome 3 is associated with lateral globus pallidus. Hits were also found for occipital fusiform gyrus (rs62621376 on chromosome 17 which is within the gene TTYH2), postcentral gyrus (rs10424191 on chromosome 19 which is located in the gene ZNF536) and planum temporale (rs5765545 on chromosome 22). In addition, three adjacent SNPs were found to be associated with the pars triangularis of inferior frontal gyrus. As for the white matter tract FA, two SNPs were identified to be associated with forceps major (rs143118835 on chromosome 7) and inferior fronto-occipital fasciculus (rs9882746 on chromosome 3). See Manhattan plots of all the global and regional measures in Fig. S1&S2.

**Fig. 3.**
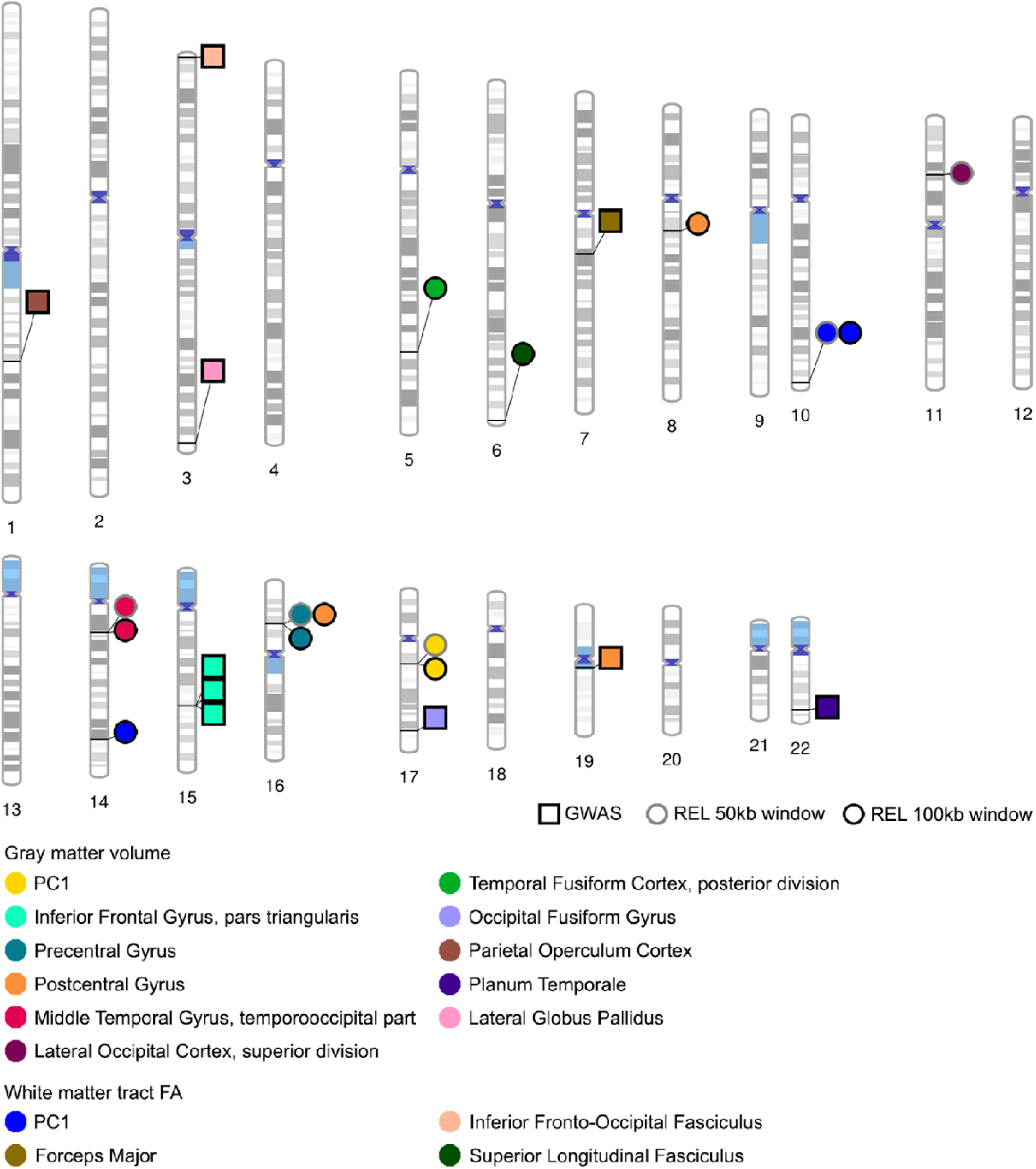
Ideogram of loci associated with global or regional brain structure measures as revealed by GWAS or REL (Figure created using PhenoGram: http://visualization.ritchielab.psu.edu/phenograms/plot). FA: fractional anisotropy.

### REL

Using the pairwise-based relatedness approach, we identified nine 100kb windows and five 50kb windows associated with either one of the two principal components or with one of the regional brain structural measures. By implication, individuals with similar traits also tend to be unusually related to each other at these particular genomic regions after correcting for background relatedness on the same chromosome. For PC1_GREY_ we found a hit on chromosome 17 where both the 50kb and 100kb windows yielded significant associations (centred at 37184.090kb and 37180.052kb separately, rs11868358 locates at the centre for both window sizes, note that loci positions were based on the reference assembly sequence GRCh37). This is to be expected because in this case the larger window only carries one more SNP compared with the smaller window (7 vs 6 SNPs). Both hit windows overlap with the gene LRRC37A11P. For PC1_WHITE_ we found two hits, one on chromosome 10, again for both the 50kb and 100kb window sizes which in this case included 22 and 35 SNPs separately (centred at 133520.594kb with the closest SNP rs11156567 and 133546.017kb with the closest SNP rs9419569 separately), the other on chromosome 14 for 100kb window size only (centred at 89578.729kb with the closest SNP rs4904507). The hit on chromosome 10 overlaps with the gene CLVS1, while the hit on chromosome 14 intersect with the gene FOXN3.

Applied to the individual measures separately, REL also revealed a number of hits, summarised in Table S4. For grey matter volume, we identified a hit on chromosome 16 showing significant association with the precentral gyrus for both 50kb and 100kb window sizes (centred at 20967.465kb and 20946.118kb with rs34051490 and rs4783513 located the closest to each window centre) with 16 and 24 SNPs included separately, overlapping with genes DNAH3, LYRM1 and DCUN1D3. Two hits with 100kb window size were found to be associated with the postcentral gyrus, one was the same hit for precentral gyrus, the other was found on chromosome 8 with 36 SNPs included (centred at 61915.362kb with rs10957177 located at the window centre). Besides, we identified a hit on chromosome 14 associated with the posterior middle temporal gyrus for both 50kb and 100kb window sizes, centred at 33903.704kb and 33877.736kb with rs11621942 and rs4982084 located at each window centre, overlapping with the gene NPAS3. For white matter tract FA, we found a 100kb window size hit associated with the superior longitudinal fasciculus on chromosome 6 (centre at 170625.624kb with 20 SNPs among which rs6923384 locates at the window entre), which intersected with genes DLL1, FAM120B, MIR4644, and LINC01624. See Manhattan plots of all the global and regional measures in Figs. S1&S2. Note that slightly more hits were found with larger windows (9 hits for 100kb and 5 hits for 50kb window size), possibly because neighbouring SNPs in high LD with the actual genetic variant(s) are more likely to be included by a larger window.

### Comparison between GWAS and REL – Empirical data

Interestingly, we observed a tendency for the two methods, GWAS and REL, to yield different results even when applied to the same dataset (see Fig. 3). To illustrate this trend we selected one trait that a hit in REL but not GWAS (PC1_WHITE_) and one trait that yielded a hit for GWAS but not REL (FA of forceps major, tract #5). For each of these two traits we then plotted p-values obtained by the two methods against each other, using a negative log scale. For both traits we analysed the data twice, once with REL ‘leading’ and once with GWAS ‘leading’. When REL was leading, we identified all windows yielding a REL p-value lower than 0.05 and plotted these against the lowest GWAS p-value among SNPs within the same window (left panel in Fig. 4 a&b). Similarly, when GWAS was leading, we identified all SNPs with a GWAS p-value lower than 0.05 and plotted these against the REL p-values of the genomic windows in which they were located (right panel in Fig. 4 a&b). As can be seen, there is little or no correlation between the sets of p-values, even for the two primary hit locations.

**Fig. 4.**
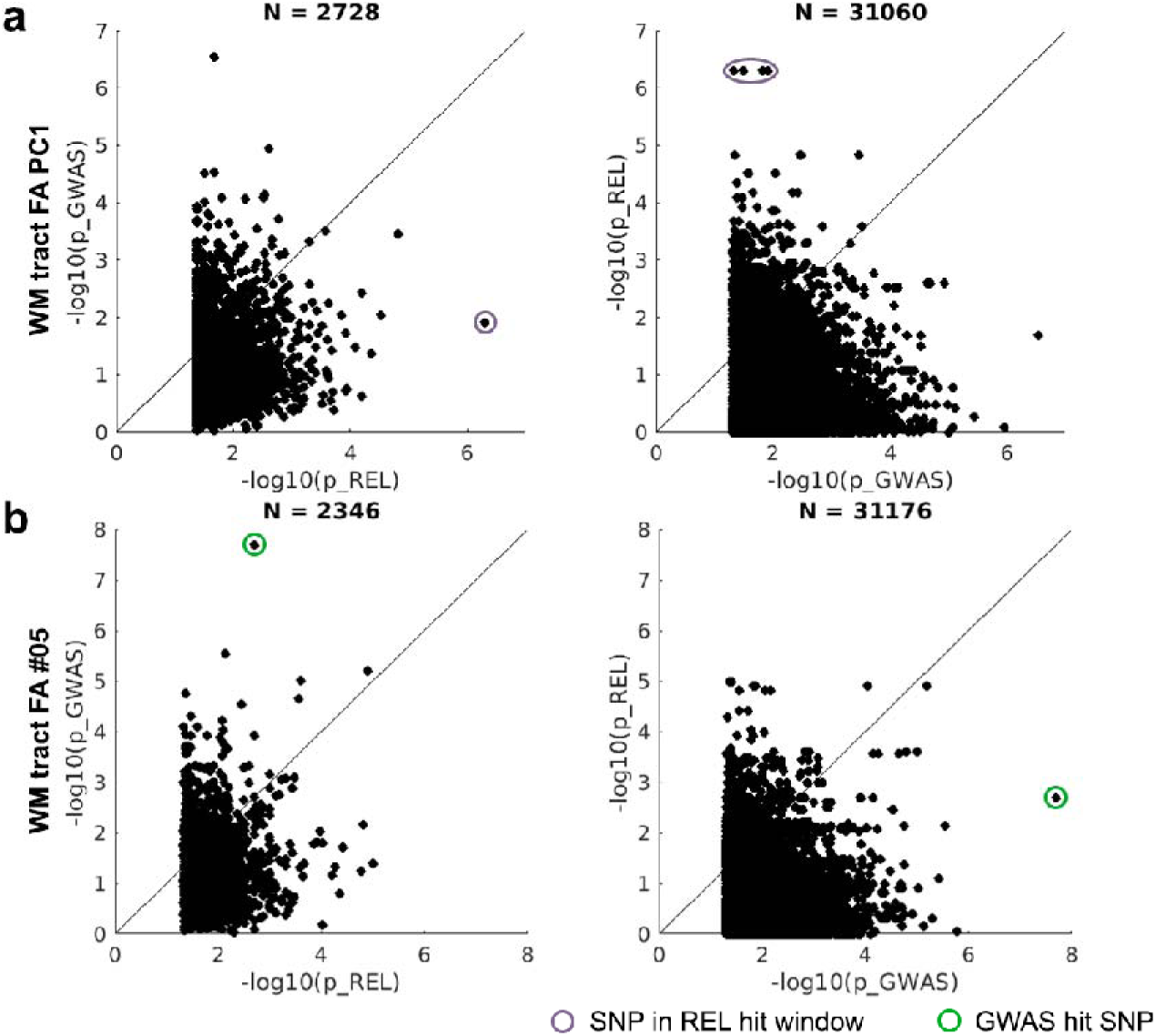
P-P plots comparing the results of GWAS and REL. In the figures on the left, 50kb genomic windows with a REL p-value less than 0.05 were first selected, their REL p-values were plotted against the lowest GWAS p-value among SNPs in the window. In the figures on the right, SNPs with a GWAS p-value less than 0.05 were selected, their GWAS p-values were plotted against the REL p-values of the genomic windows they belong to. Note that REL p-value was obtained from permutation test. GWAS p-value is the Wald p-value. (a) Results of the first principle component of white matter tract FA where a hit window was identified in REL. (b) Results of the white matter tract #05 where a GWAS hit was found. N: number of genomic windows (left) or SNPs (right) above threshold.

One important difference between GWAS and REL is that GWAS is based on individual SNPs, making it potentially susceptible to undue influence by outliers, while relatedness in REL is based on multiple observations, making it less likely to be affected by outliers. This difference is illustrated in Fig. 5a, where we compare a standard GWAS analysis with a GWAS analysis based on robust regression. In several instances, the original GWAS hit is clearly driven by just one or two extreme data points and significance is lost when robust regression is applied (see the results of GM volume #52 and WM tract FA #7). In contrast, reanalysing the REL hits with and without robust regression reveals very little impact on significance for both the 50kb and 100kb window sizes alike (Fig. 5b).

**Fig. 5.**
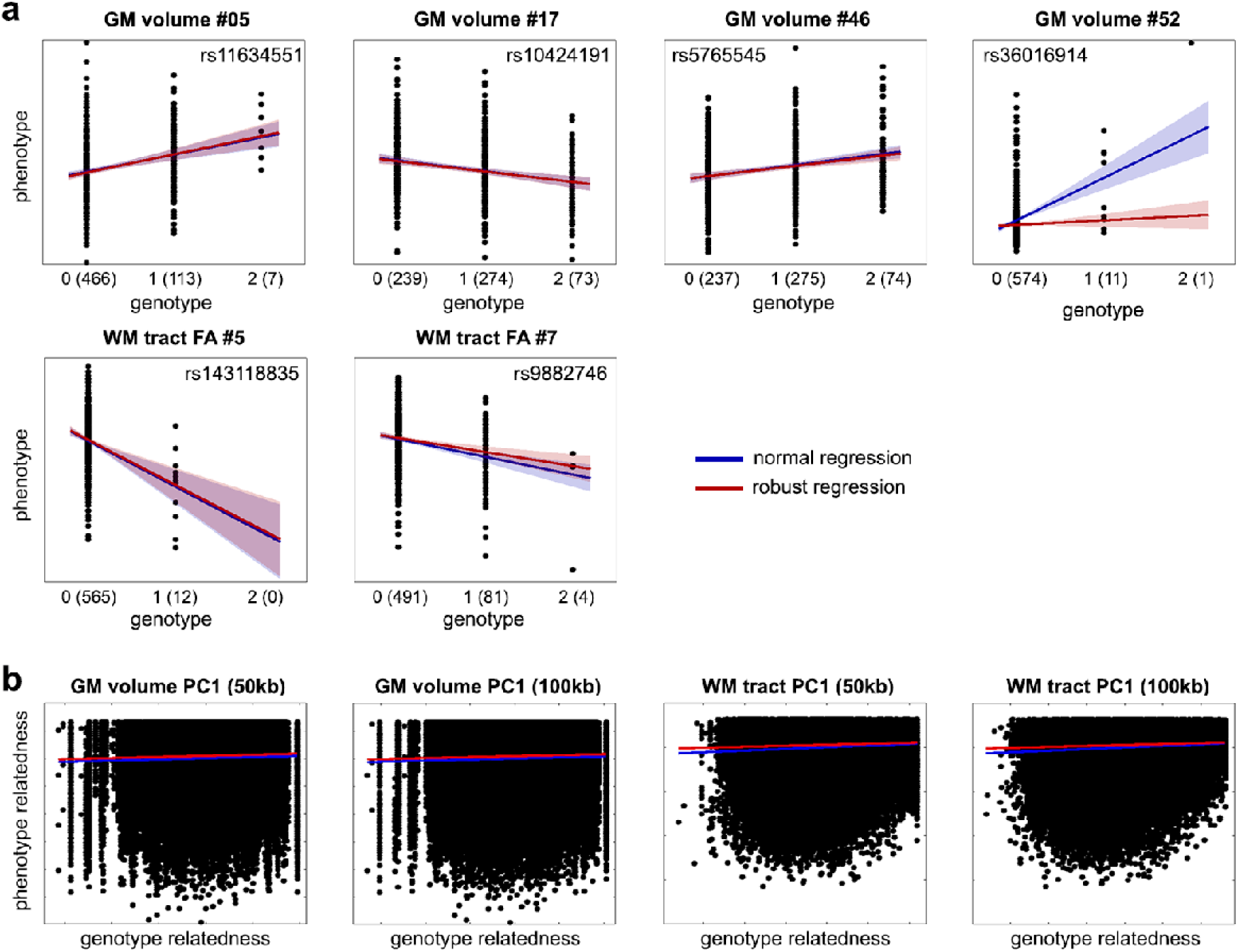
GWAS and REL hits with or without robust regression. (a) Loci associated with grey matter and white matter measures identified in GWAS. Number of individuals is listed in the parentheses after each additively coded genotype along the x-axis (0 - homozygous reference alleles, 1 - heterozygous alleles, 2 - homozygous alternative alleles). (b) REL results of the first principle component of grey matter volume and white matter tract FA for both 50kb- and 100kb window. Shade indicates 95% confidence interval.

### Comparison between GWAS and REL – Simulation

To compare GWAS with REL directly, we conducted a simulation analysis to evaluate their performance with respect to local levels of LD. We selected 100 genomic windows covering a wide range of LD values. Within each window we selected one SNP to act as the ‘functional’ SNP associated with the trait of interest. To create this association, we re-assigned trait values according to the genotype of the selected functional SNP. In this way we preserve the characteristics of the real data in terms of LD between SNPs within the window. The simulated signal injected into a genomic window can be maximized by selecting as the functional SNP the SNP in the highest overall LD with the other SNPs in the window. Similarly, signal strength can be reduced by selecting the functional SNP to be the one with lowest LD to other SNPs. A second way to modulate signal strength is to add Gaussian noise to the trait values.

As shown in Fig. 6a, even for a relatively low signal to noise ratio (SNR), windows with high average LD (i.e., mean pair-wise LD among SNPs within a window) still reached the minimal p-value (i.e., 0.001 given 1,000 permutations) in REL. GWAS also successfully identified simulated functional SNPs when these were in high LD with other SNPs in the window. It can be seen that SNPs in positive LD with the simulated functional SNP are distinguished from SNPs that are in negative LD, an effect that gets stronger as the SNR is increased. The same trend can also be found when the SNP that has the lowest overall LD was selected as the functional SNP, but it requires a higher SNR to achieve similar patterns (Fig. 6b).

**Fig. 6.**
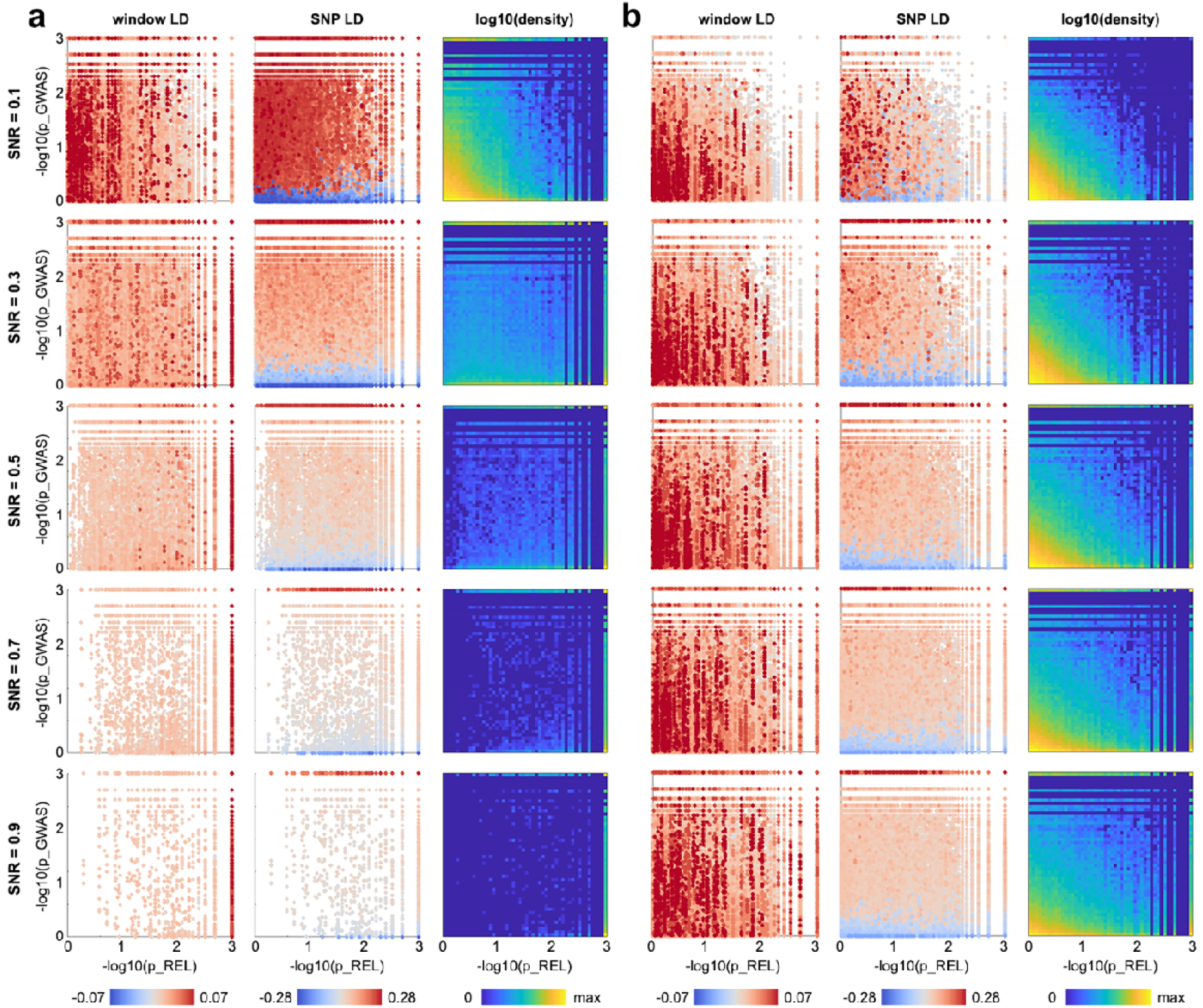
Simulation results for GWAS and REL with different SNRs. Both GWAS p-value and REL p-value were based on non-parametric permutation test. Dots are colored by the LD of genomic window or SNP, or the density calculated as the percentage of dots in each 0.6 x 0.6 grid. (a) Simulation results with the SNP in the highest overall LD with the other SNPs in the genomic window as functional SNP. (b) Simulation results with the SNP in the lowest overall LD with the other SNPs as functional SNP.

Finally, we compared the performance of the two methods by calculating the ratio of dots in the upper left triangle to those in the lower right triangle in the p-p plots in Fig. 7. As shown in Fig. 7a, GWAS is more sensitive when the signal is very weak, but it is outperformed by REL as the SNR increases. We also calculated the detection rate of GWAS and REL, which is defined as the ratio of cases where the p-value reaches the most significant level given the number of permutations (i.e., 1/1001 for 1000 permutations). In terms of the detection rate, GWAS also outperformed REL when the signal is relatively weak, while REL outperformed GWAS when the SNR increases, especially when the functional SNP was selected as the one in high LD with the other SNPs within the genomic window (Fig. 7 b&c).

**Fig. 7.**
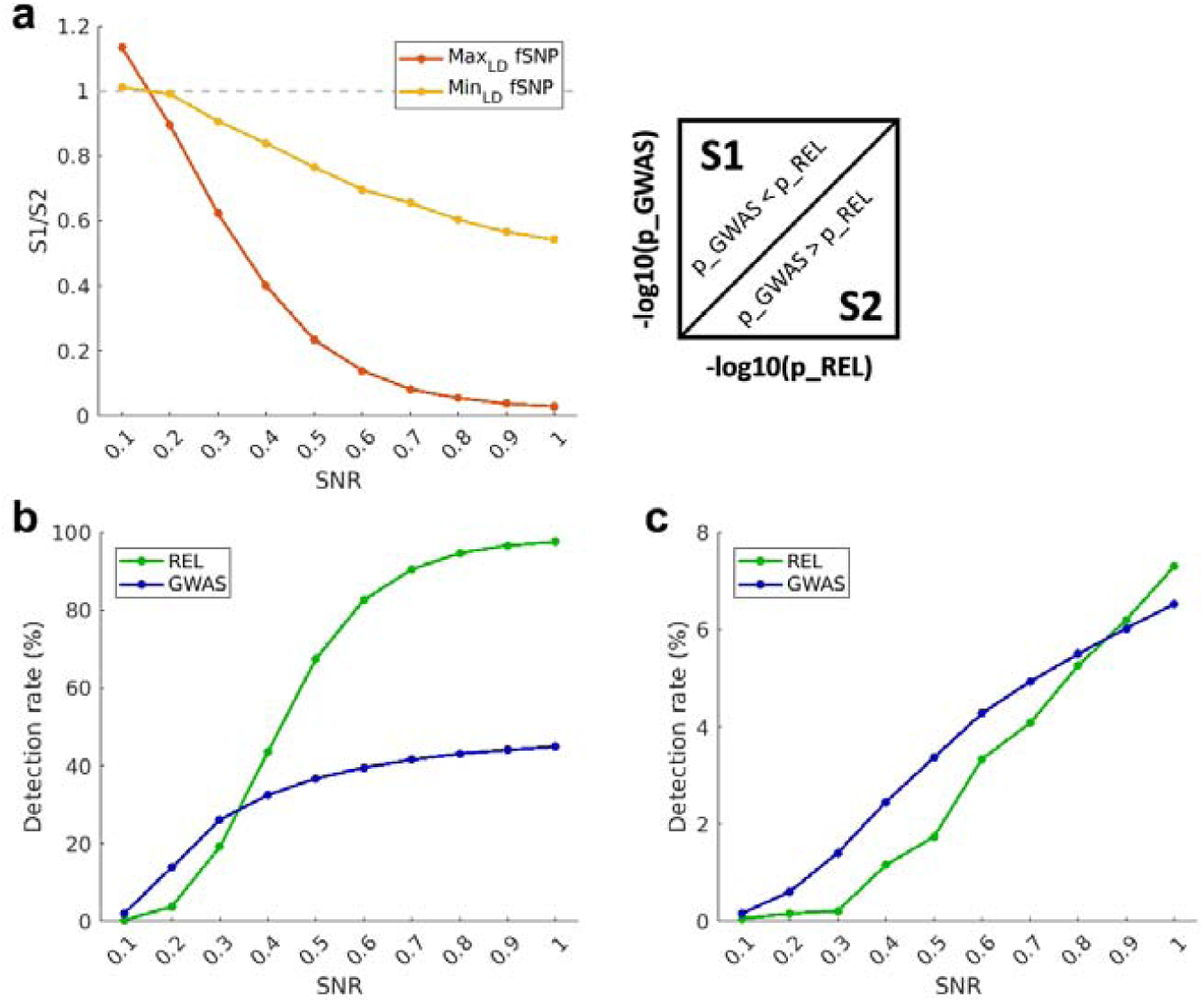
Summary of the performance of GWAS and REL for detecting simulated signals with different SNRs. (a) Ratio of the cases where GWAS outperforms REL to the cases the other way around. S1: number of dots in the upper left triangle. S2: number of dots in the lower right triangle. Max_LD_ fSNP: functional SNP was selected as the one in highest overall LD with the other SNPs. Min_LD_ fSNP: functional SNP was selected as the one in lowest overall LD with the other SNPs. (b) Detection rate (percentage of cases where permutation p-value reached the minimal value, i.e., no outliers found in 1000 permutations) for Max_LD_ fSNP. (c) Detection rate for Min_LD_ fSNP.

## Discussion

In this study we compared the performance of a classical GWAS analysis with REL, an alternative, regression-based approach involving pairwise comparisons between individuals, that allows genetic relatedness to be included in the model. We applied both approaches to a large data set of grey and while matter brain measurements. Classical GWAS is potentially very powerful if the functional SNP has been sampled, but is likely to be much weaker when there is reliance on linkage disequilibrium between a sampled SNP and an unsampled functional SNP. Wherever GWAS tests each SNP separately, the large number of tests required reduces power because of the need to correct for relatively higher rates of false positives. In comparison, the regression-based analysis may offer higher power by integrating information across multiple SNPs within a genomic window, thereby greatly reducing the number of tests. By focusing on relatedness, the regression method may be efficient at picking up signals when the functional SNP has not been sampled, though is likely to be less powerful than a GWAS when the functional SNP has been sampled.

We began by summarising the measurements in the sample using PCA and testing the first two PCs as well as individual ROIs of both grey matter and white matter. Both approaches gave a similar number of hits (10 hits in GWAS, 9 hits with 100kb window size and 5 hits with 50kb window size in REL). However, some GWAS hits appeared to be driven by outliers and were lost when robust regression was applied. Our use of full Bonferroni correction is conservative, but we feel this is appropriate given the extent to which small numbers of outliers are able to impact the analysis, particularly the GWAS. Moreover, a genetic association is possibly driven by more than one variant, perhaps suggested by the way neighbouring SNPs give opposing trends (see the middle column in Fig. 6a&b). It is a further potential advantage of the regression approach that it is able, under some circumstances, to integrate over multiple variants.

Among the loci we identified in GWAS and REL (see Tables S1&S2), the genomic window associated with the overall grey matter structure (i.e., PC1_grey_) overlapped with LRRC37A11P which belongs to a family of genes expressed during human corticogenesis ^30^. The genomic window influencing the volume of the posterior middle temporal gyrus overlapped with the gene NPAS3 which has an important role in brain development ^31, 32, 33^, it is also associated with learning disability and mental disorders such as schizophrenia ^34, 35^. The genomic window associated with the FA of the superior longitudinal fasciculus covered genes DLL1, FAM120B and MIR4644, these neighboring genes have been reported to be responsible for structural brain abnormalities with a hypothesized leading role of DLL1 in interacting with other adjacent genes ^36, 37, 38, 39, 40^. Moreover, DLL1 has also been identified as a candidate gene for intellectual disability ^39^. The REL hit related to the volume of postcentral gyrus covered the gene DNAH3 which has significant associations with major depressive disorder ^41^. While the GWAS hit for the same brain phenotype was found within the gene ZNF536 which is highly expressed in the developing central nervous system and is related to neural differentiation ^42^. Moreover, some genes identified in our study were reported to be associated with brain structure (e.g., PCDHGA1 ^43^), healthy aging (e.g., PCDHGA3 ^44^) and intractable epilepsy (e.g., SLC35A2 ^45^).

We were interested in testing the possibility that relatedness between individuals at or near a functional variant provides a more generally applicable indicator that trait values are similar compared with a classical GWAS. For this, we conducted a simulation analysis in which we chose a selection of windows covering a wide range of levels of LD. Within each window we created a correlation between phenotype and genotype at one SNP, removed this SNP, and then tested to see whether local LD is sufficient to drive an association. In practice, higher LD with the functional SNP resulted in better performance in both GWAS and REL. GWAS is more sensitive than REL when the simulated signal is weak, while REL tends to outperform GWAS as the strength of simulated signal increases. This analysis suggests that the best approach may vary across the genome and that pre-testing for LD could allow GWAS to be used where LD is high and regression to be used where LD is low, or the local pattern of LD is complex.

Age-related effects typically introduce large sources of variance in studies of humans, including the current study. All of our brain phenotype measures vary with age, often strongly so. This raises questions about how best to achieve control. Here we have used a simple linear correction. While this appears to be generally effective, it may be possible to improve the method by using a non-linear correction. Equally, although some form of correction is undoubtedly necessary, a simple linear correction has the potential to introduce age-distribution biases wherever appreciably deviations from a linear relationship exist. Conversely, it is possible to imagine genetic factors that operate through either emphasising or ameliorating age-related changes, and these could be lost through over-aggressive control. We hope that a more in-depth analysis of how best to control for age-related effects will be the subject of future studies.

Association studies have enjoyed a rather chequered history. Early studies were based on modest sample sizes but also modest marker numbers. Progression from microsatellites to SNPs saw a dramatic increase in marker number, from a few hundreds to several hundreds of thousands, an increase that was accompanied by concerns about the rate of false positives ^46, 47, 48, 49, 50^. To compensate, many studies aimed to increase sample sizes to thousands, tens of thousands or more. In addition, the statistical methods have been constantly refined, while genome sequencing coupled with imputation has further increased marker coverage. In comparison, our study uses modest sample sizes. However, our dataset is like a number of others in being limited by extrinsic factors, in our case by the number of individuals for which in-depth phenotypic data could be collected. Similarly, we deliberately avoided the most recent ‘cutting edge’ GWAS methods so that our results are broadly comparable to a range of prior studies. Despite this, we still find a number of reasonably convincing hits that lead us to believe that our approach can offer a novel view both for other smaller datasets ^51^ and, with development, as a method that could offer improvements to the analysis of larger datasets as well.

In conclusion, we have systematically compared the classical GWAS with a pair-wise relatedness-based approach, REL, using empirical data from human brain structural measures, as well as simulated data which evaluated the performance of these two approaches with respect to local LD and signal strength. Both approaches succeeded in identifying genetic variants associated with brain structure, though they tend to give distinct results given the very different aspects of genetic properties used. Moreover, simulation results show that GWAS outperforms REL when the signal is relatively weak. As the signal strength increases, the collective effects due to local LD boost the performance of REL, making it more successful than GWAS in picking up the signal. Thus, REL provides an alternative way to provide complementary results for the search of genetic associations in future studies.

## Acknowledgments

We thank Prof. Simon E. Fisher and Dr. Else Eising for processing the raw genotypes and comments on the manuscript. This work was supported by a European Research Council Advanced Investigator Grant to L.K.T. under the European Community’s Horizon 2020 Research and Innovation Programme (2014–2020 ERC grant agreement number 669820) and a European Union Horizon 2020 Grant: ‘Healthy minds 0–100 years: Optimising the use of European brain imaging cohorts (“Lifebrain”) to L.K.T. (grant agreement number 732592). K.A.T. was supported by the British Academy (PF160048) and the Guarantors of Brain (G101149). The Cambridge Centre for Aging and Neuroscience (Cam-CAN) research was supported by the Biotechnology and Biological Sciences Research Council (grant number BB/H008217/1).

## REFERENCES

1. Honey CJ, Thivierge J-P, Sporns O. Can structure predict function in the human brain? Neuroimage 52, 766–776 (2010).

2. Webb CE, Rodrigue KM, Hoagey DA, Foster CM, Kennedy KM. Contributions of White Matter Connectivity and BOLD Modulation to Cognitive Aging: A Lifespan Structure-Function Association Study. Cereb Cortex 30, 1649–1661 (2019).

3. Zatorre RJ, Fields RD, Johansen-Berg H. Plasticity in gray and white: neuroimaging changes in brain structure during learning. Nature Neuroscience 15, 528–536 (2012).

4. Andreasen N, et al. Intelligence and brain structure in normal individuals. The American journal of psychiatry 150, 130–134 (1993).

5. Chen P-Y, Chen C-L, Hsu Y-C, Tseng W-YI. Fluid intelligence is associated with cortical volume and white matter tract integrity within multiple-demand system across adult lifespan. Neuroimage 212, 116576 (2020).

6. Gazes Y, et al. fMRI-guided white matter connectivity in fluid and crystallized cognitive abilities in healthy adults. Neuroimage 215, 116809 (2020).

7. Hibar DP, et al. Novel genetic loci associated with hippocampal volume. Nature Communications 8, 13624 (2017).

8. Hibar DP, et al. Common genetic variants influence human subcortical brain structures. Nature 520, 224 (2015).

9. Hofer E, et al. Genetic Determinants of Cortical Structure (Thickness, Surface Area and Volumes) among Disease Free Adults in the CHARGE Consortium. bioRxiv, 409649 (2019).

10. Biton A, et al. Polygenic architecture of human neuroanatomical diversity. bioRxiv, 592337 (2019).

11. Grasby KL, et al. The genetic architecture of the human cerebral cortex. Science 367, eaay6690 (2020).

12. Rodrigue AL, et al. Evidence for genetic correlation between human cerebral white matter microstructure and inflammation. Human Brain Mapping 0, (2019).

13. Raj SM, Pagani L, Gallego Romero I, Kivisild T, Amos W. A general linear model-based approach for inferring selection to climate. BMC Genetics 14, 87 (2013).

14. Kriegeskorte N, Mur M, Bandettini PA. Representational similarity analysis-connecting the branches of systems neuroscience. Frontiers in systems neuroscience 2, 4 (2008).

15. Reich DE, et al. Linkage disequilibrium in the human genome. Nature 411, 199–204 (2001).

16. Li M-X, Gui H, Kwan JSH, Sham PC. GATES: a rapid and powerful gene-based association test using extended Simes procedure. Am J Hum Genet 88 3, 283–293 (2011).

17. Ehret GB, Lamparter D, Hoggart CJ, Whittaker JC, Beckmann JS, Kutalik Zn. A multi-SNP locus-association method reveals a substantial fraction of the missing heritability. Am J Hum Genet 91 5, 863–871 (2012).

18. Zhao H, Nyholt DR, Yang Y, Wang J, Yang Y. Improving the detection of pathways in genome-wide association studies by combined effects of SNPs from Linkage Disequilibrium blocks. Scientific Reports 7, (2017).

19. de Leeuw CA, Mooij JM, Heskes T, Posthuma D. MAGMA: Generalized Gene-Set Analysis of GWAS Data. PLoS Comput Biol 11, e1004219 (2015).

20. Sul JH, Martin LS, Eskin E. Population structure in genetic studies: Confounding factors and mixed models. PLOS Genetics 14, e1007309 (2018).

21. Vilhjálmsson BJ, Nordborg M. The nature of confounding in genome-wide association studies. Nature Reviews Genetics 14, 1–2 (2013).

22. Shafto MA, et al. The Cambridge Centre for Ageing and Neuroscience (Cam-CAN) study protocol: a cross-sectional, lifespan, multidisciplinary examination of healthy cognitive ageing. BMC Neurology 14, 204 (2014).

23. Guo J, et al. Resin-assisted enrichment of thiols as a general strategy for proteomic profiling of cysteine-based reversible modifications. Nature protocols 9, 64 (2014).

24. Purcell S, et al. PLINK: a tool set for whole-genome association and population-based linkage analyses. Am J Hum Genet 81, 559–575 (2007).

25. Taylor JR, et al. The Cambridge Centre for Ageing and Neuroscience (Cam-CAN) data repository: Structural and functional MRI, MEG, and cognitive data from a cross-sectional adult lifespan sample. Neuroimage 144, 262–269 (2017).

26. Pierpaoli C, Jezzard P, Basser PJ, Barnett A, Di Chiro G. Diffusion tensor MR imaging of the human brain. Radiology 201, 637–648 (1996).

27. Zhou X, Stephens M. Genome-wide efficient mixed-model analysis for association studies. Nature genetics 44, 821–824 (2012).

28. Queller DC, Goodnight KF. Estimating relatedness using genetic markers. Evolution 43, 258–275 (1989).

29. Ragsdale AP, Gravel S. Unbiased Estimation of Linkage Disequilibrium from Unphased Data. Molecular Biology and Evolution, (2019).

30. Suzuki IK, et al. Human-Specific NOTCH2NL Genes Expand Cortical Neurogenesis through Delta/Notch Regulation. Cell 173, 1370-1384.e1316 (2018).

31. Kamm GB, Pisciottano F, Kliger R, Franchini LF. The developmental brain gene NPAS3 contains the largest number of accelerated regulatory sequences in the human genome. Molecular biology and evolution 30, 1088–1102 (2013).

32. Gould P, Kamnasaran D. Immunohistochemical analyses of NPAS3 expression in the developing human fetal brain. Anatomia, histologia, embryologia 40, 196–203 (2011).

33. Sha L, et al. Transcriptional regulation of neurodevelopmental and metabolic pathways by NPAS3. Molecular psychiatry 17, 267–279 (2012).

34. Pickard BS, Malloy M, Porteous D, Blackwood D, Muir W. Disruption of a brain transcription factor, NPAS3, is associated with schizophrenia and learning disability. American Journal of Medical Genetics Part B: Neuropsychiatric Genetics 136, 26–32 (2005).

35. Pickard BS, Pieper AA, Porteous DJ, Blackwood DH, Muir WJ. The NPAS3 gene—emerging evidence for a role in psychiatric illness. Annals of medicine 38, 439–448 (2006).

36. Kawaguchi D, Furutachi S, Kawai H, Hozumi K, Gotoh Y. Dll1 maintains quiescence of adult neural stem cells and segregates asymmetrically during mitosis. Nature communications 4, 1–12 (2013).

37. Kim W-Y, Shen J. Presenilins are required for maintenance of neural stem cells in the developing brain. Molecular neurodegeneration 3, 2 (2008).

38. Peddibhotla S, et al. Delineation of candidate genes responsible for structural brain abnormalities in patients with terminal deletions of chromosome 6q27. European Journal of Human Genetics 23, 54–60 (2015).

39. Hanna MD, et al. Defining the Critical Region for Intellectual Disability and Brain Malformations in 6q27 Microdeletions. Molecular syndromology 10, 202–208 (2019).

40. Fischer-Zirnsak B, et al. Haploinsufficiency of the Notch Ligand DLL1 Causes Variable Neurodevelopmental Disorders. The American Journal of Human Genetics 105, 631–639 (2019).

41. Jawinski P, et al. Human brain arousal in the resting state: a genome-wide association study. Molecular psychiatry 24, 1599–1609 (2019).

42. Qin Z, et al. ZNF536, a novel zinc finger protein specifically expressed in the brain, negatively regulates neuron differentiation by repressing retinoic acid-induced gene transcription. Molecular and cellular biology 29, 3633–3643 (2009).

43. Zhao B, et al. Transcriptome-wide association analysis of 211 neuroimaging traits identifies new genes for brain structures and yields insights into the gene-level pleiotropy with other complex traits. bioRxiv, 842872 (2019).

44. Kim S, et al. DNA methylation associated with healthy aging of elderly twins. Geroscience 40, 469–484 (2018).

45. Sim NS, et al. Brain somatic mutations in SLC35A2 cause intractable epilepsy with aberrant N-glycosylation. Neurology Genetics 4, e294 (2018).

46. Shafquat A, Crystal RG, Mezey JG. Identifying novel associations in GWAS by hierarchical Bayesian latent variable detection of differentially misclassified phenotypes. BMC Bioinformatics 21, 178 (2020).

47. Visscher PM, et al. 10 Years of GWAS Discovery: Biology, Function, and Translation. The American Journal of Human Genetics 101, 5–22 (2017).

48. Tam V, Patel N, Turcotte M, Bossé Y, Paré G, Meyre D. Benefits and limitations of genome-wide association studies. Nature Reviews Genetics 20, 467–484 (2019).

49. Korte A, Farlow A. The advantages and limitations of trait analysis with GWAS: a review. Plant Methods 9, 29 (2013).

50. Fadista J, Manning AK, Florez JC, Groop L. The (in)famous GWAS P-value threshold revisited and updated for low-frequency variants. European Journal of Human Genetics 24, 1202–1205 (2016).

51. Quick C, et al. Sequencing and imputation in GWAS: Cost-effective strategies to increase power and genomic coverage across diverse populations. Genetic Epidemiology n/a, (2020).

