## Supplemental Figures 1-2 for "Genetic signatures of human brain structure: A comparison between GWAS and relatedness-based regression"

**Figure S1** GWAS and REL results of grey matter volume (PC1, PC2 and 55 ROIs). Left: GWAS results (Wald p-value), middle & right: REL results of 50kb & 100kb genomic windows (Permutation p-value). Red horizontal bar: genome-wide Bonferroni correction threshold. Yellow horizontal bar: chromosome-wide Bonferroni correction threshold.

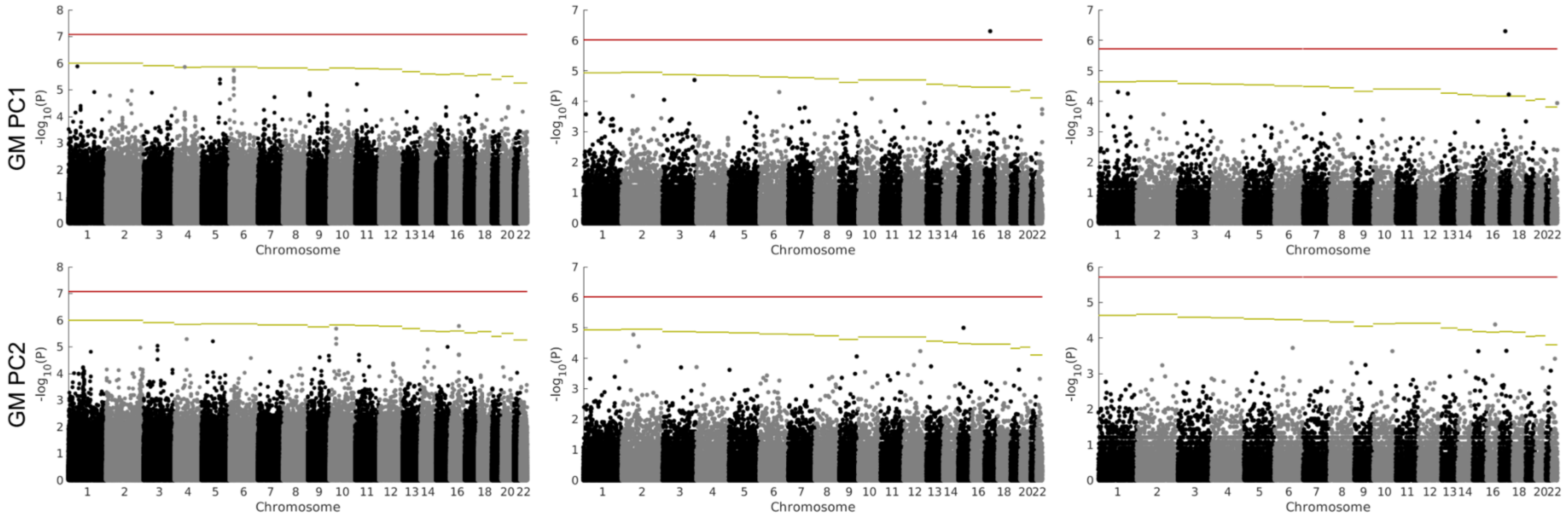

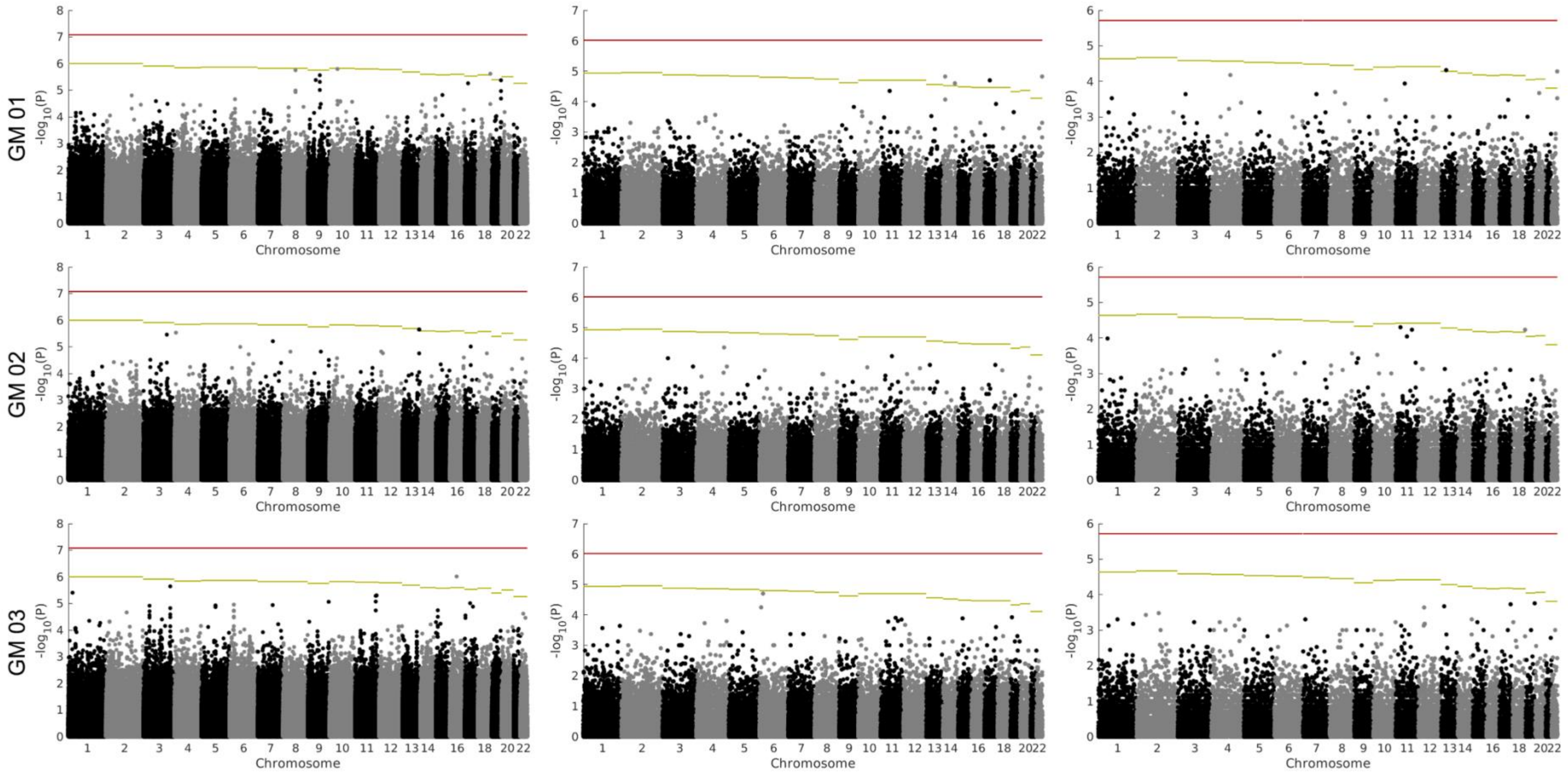
GM 01
GM 02
GM 03
-log10(P)
Chromosome
1 2 3 4 5 6 7 8 9 10 11 12 13 14 16 18 20 22

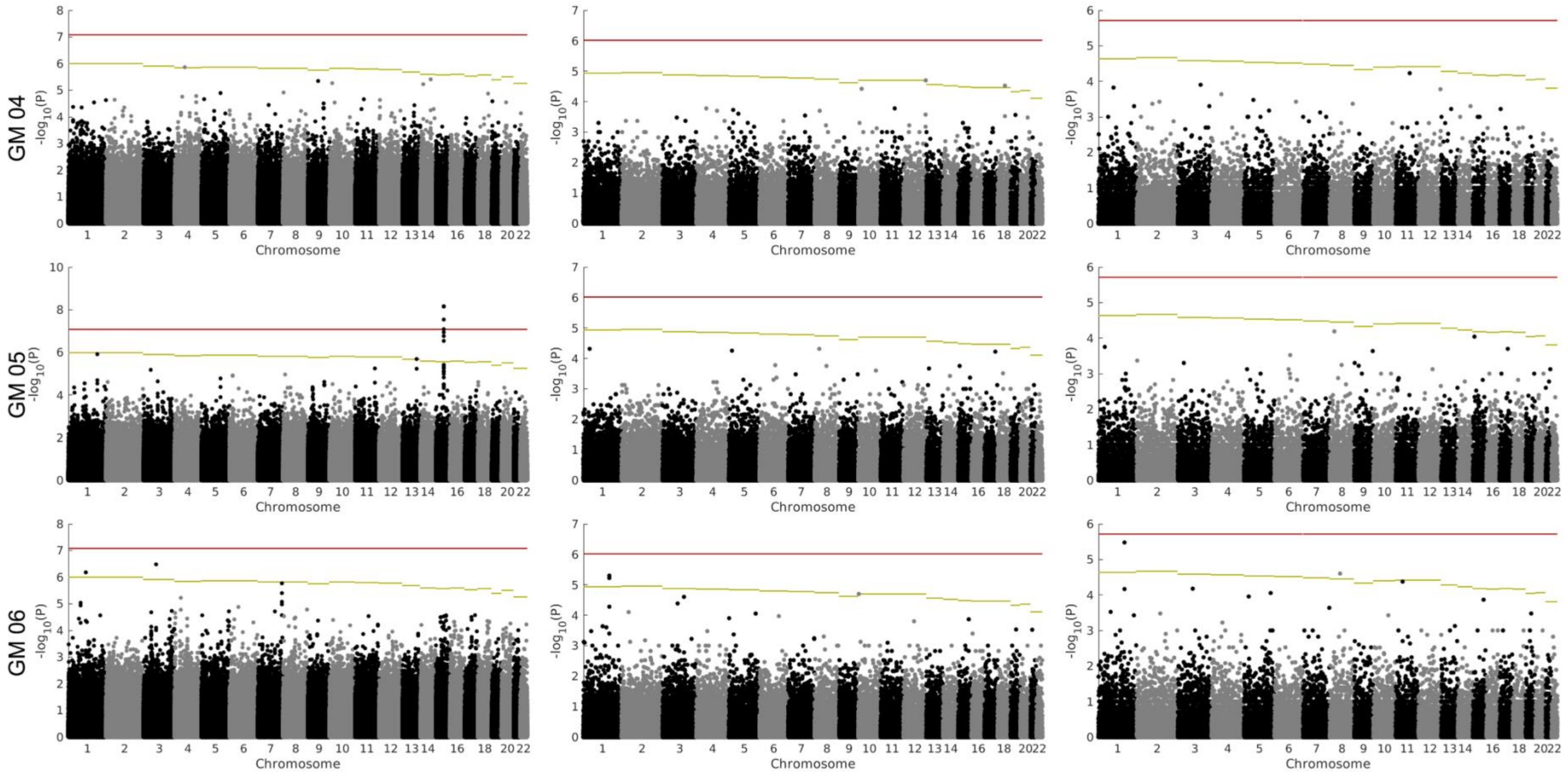
GM 04
GM 05
GM 06
-log10(P)
Chromosome

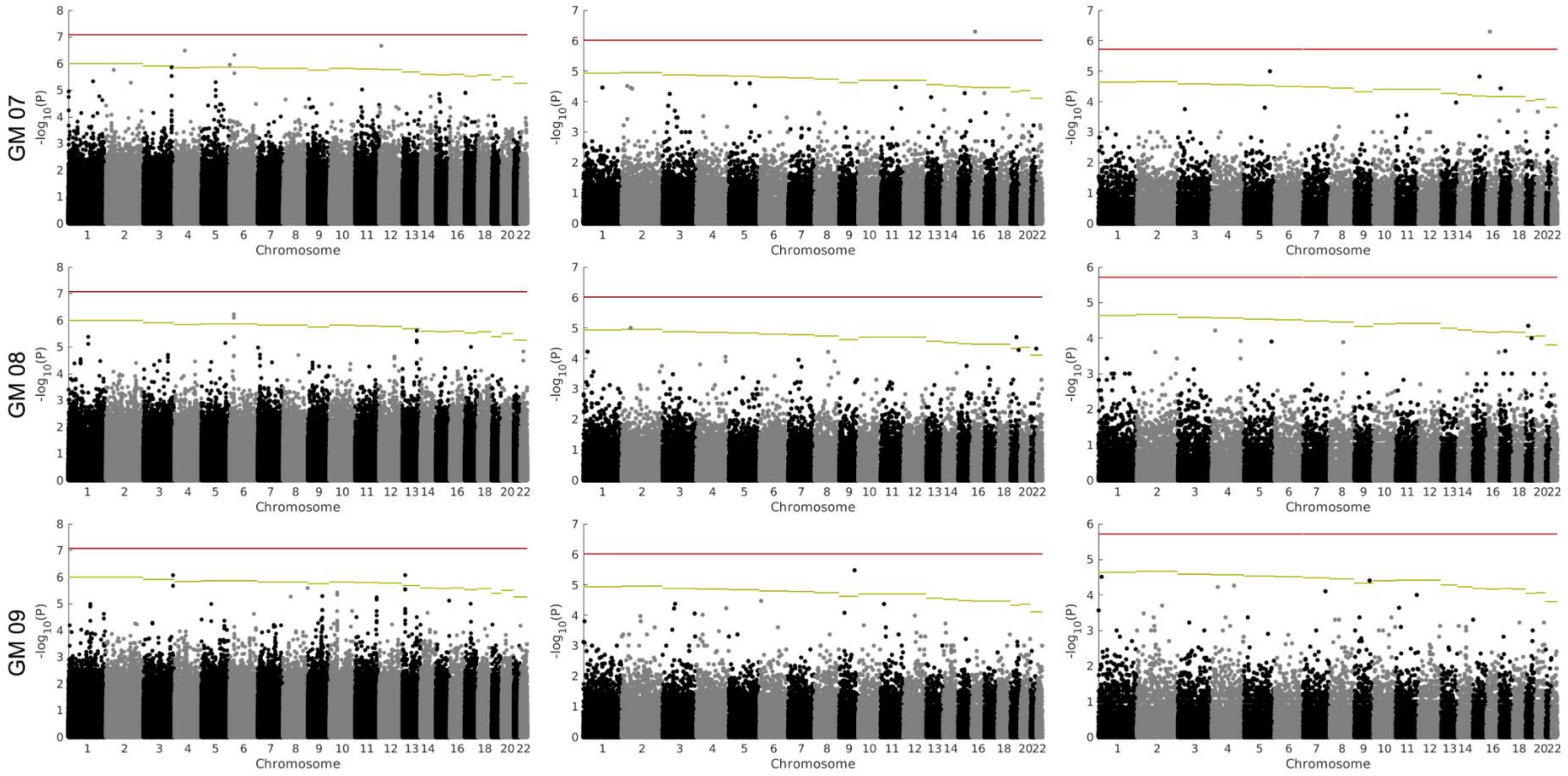
GM 07
GM 08
GM 09
-log10(P)
Chromosome

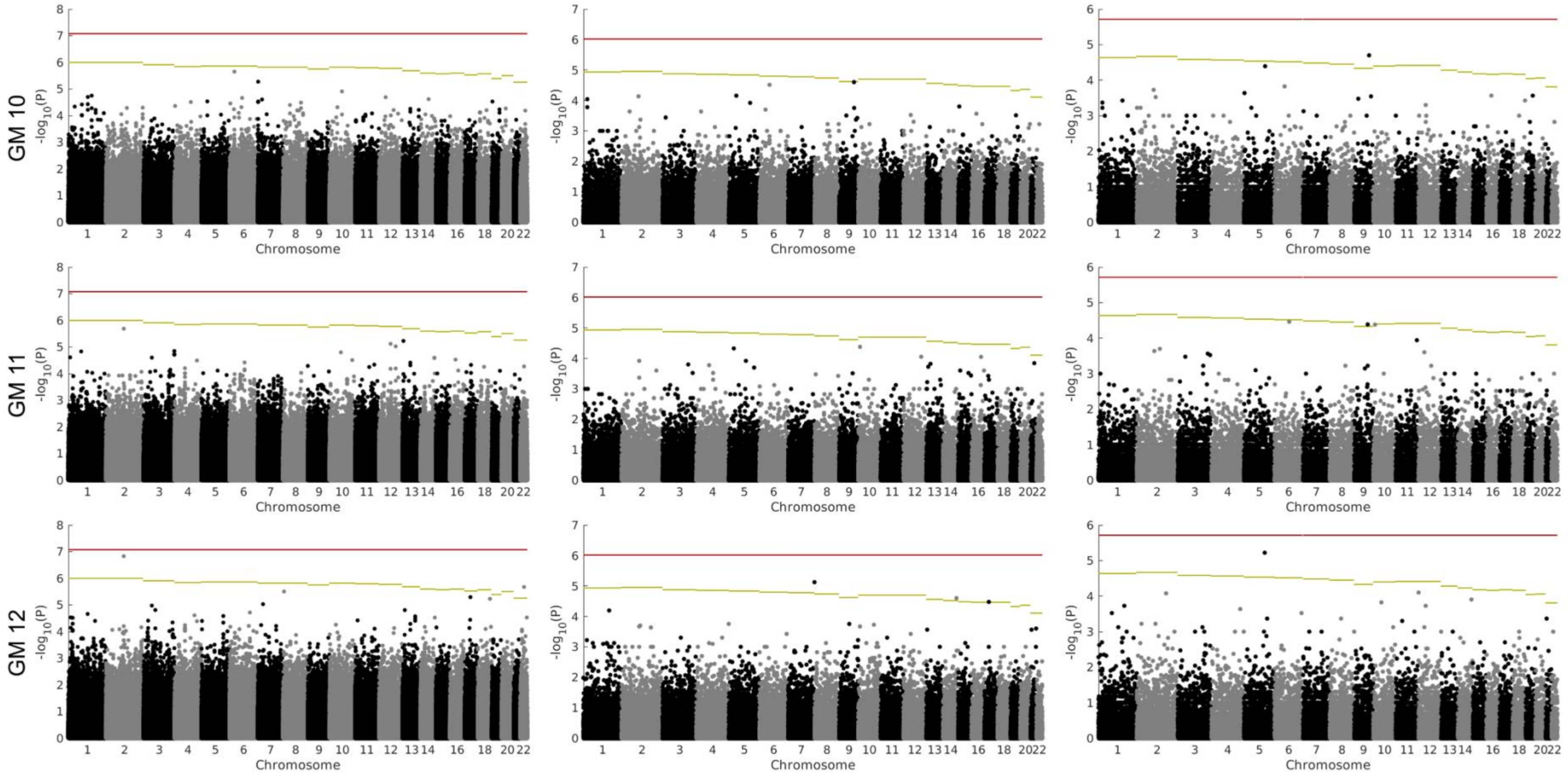

GM 10
GM 11
GM 12
-log10(P)
Chromosome

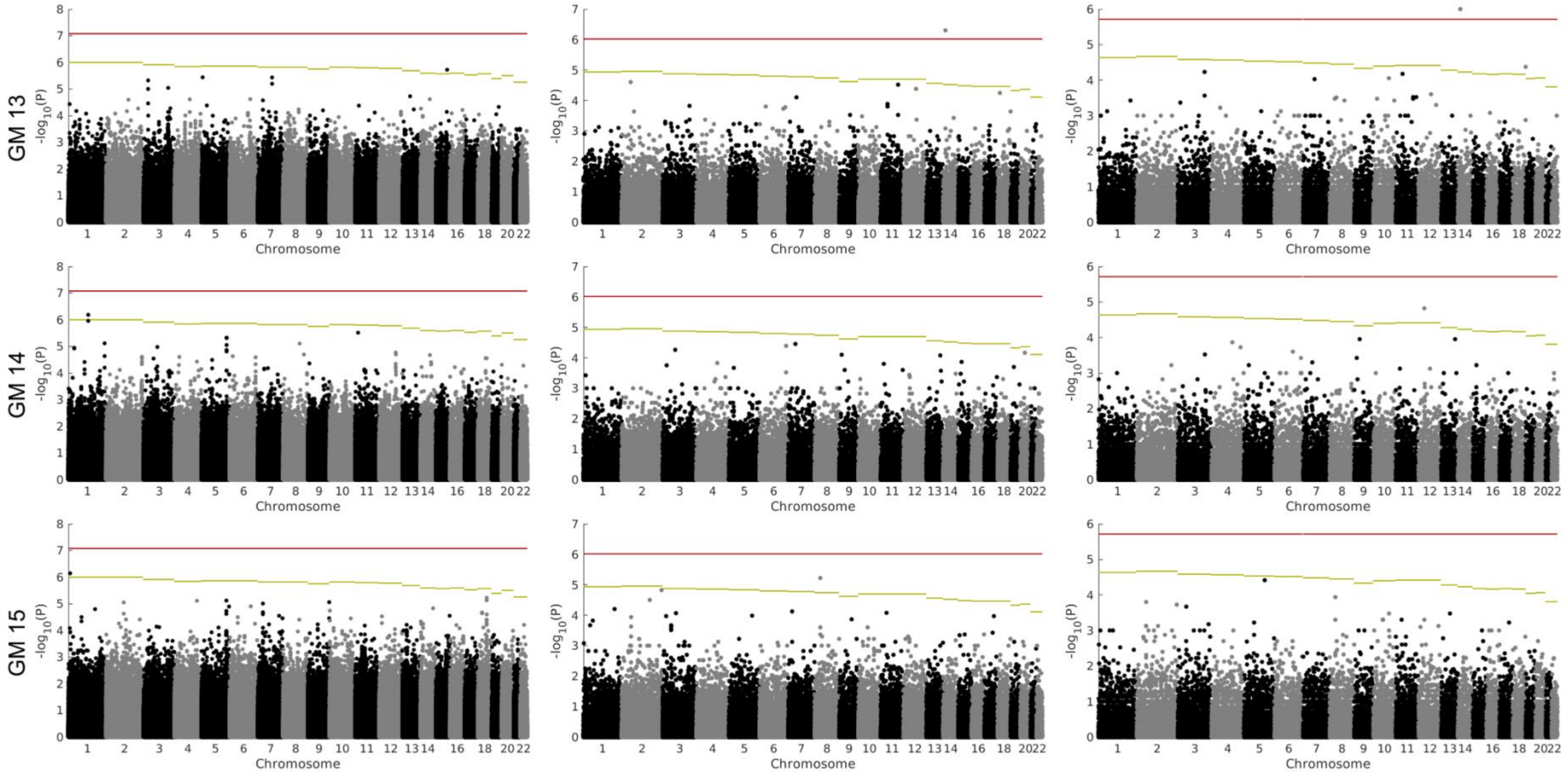

GM 13
GM 14
GM 15
-log10(P)
Chromosome

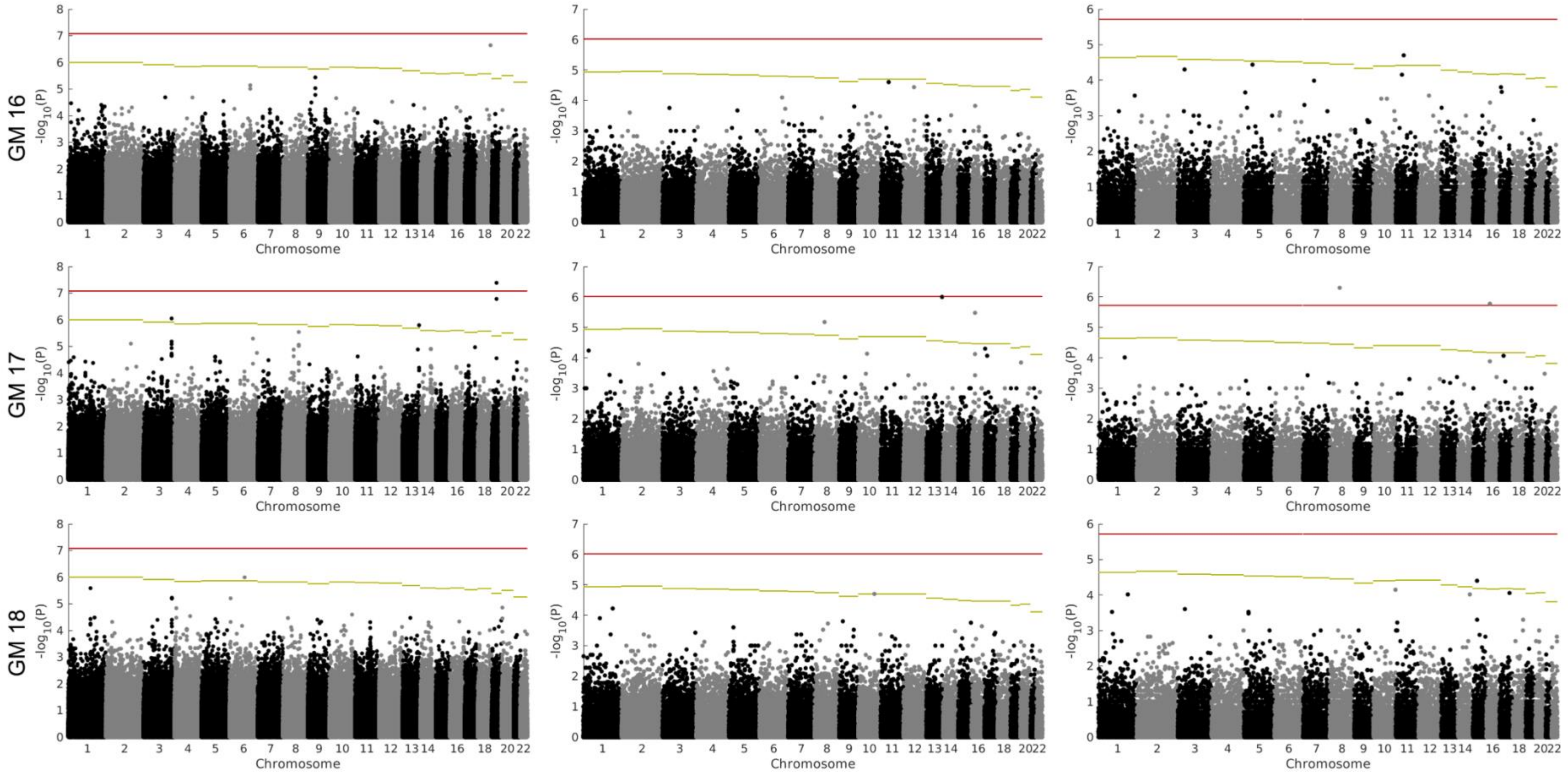
GM 16
GM 17
GM 18
-log10(P)
Chromosome

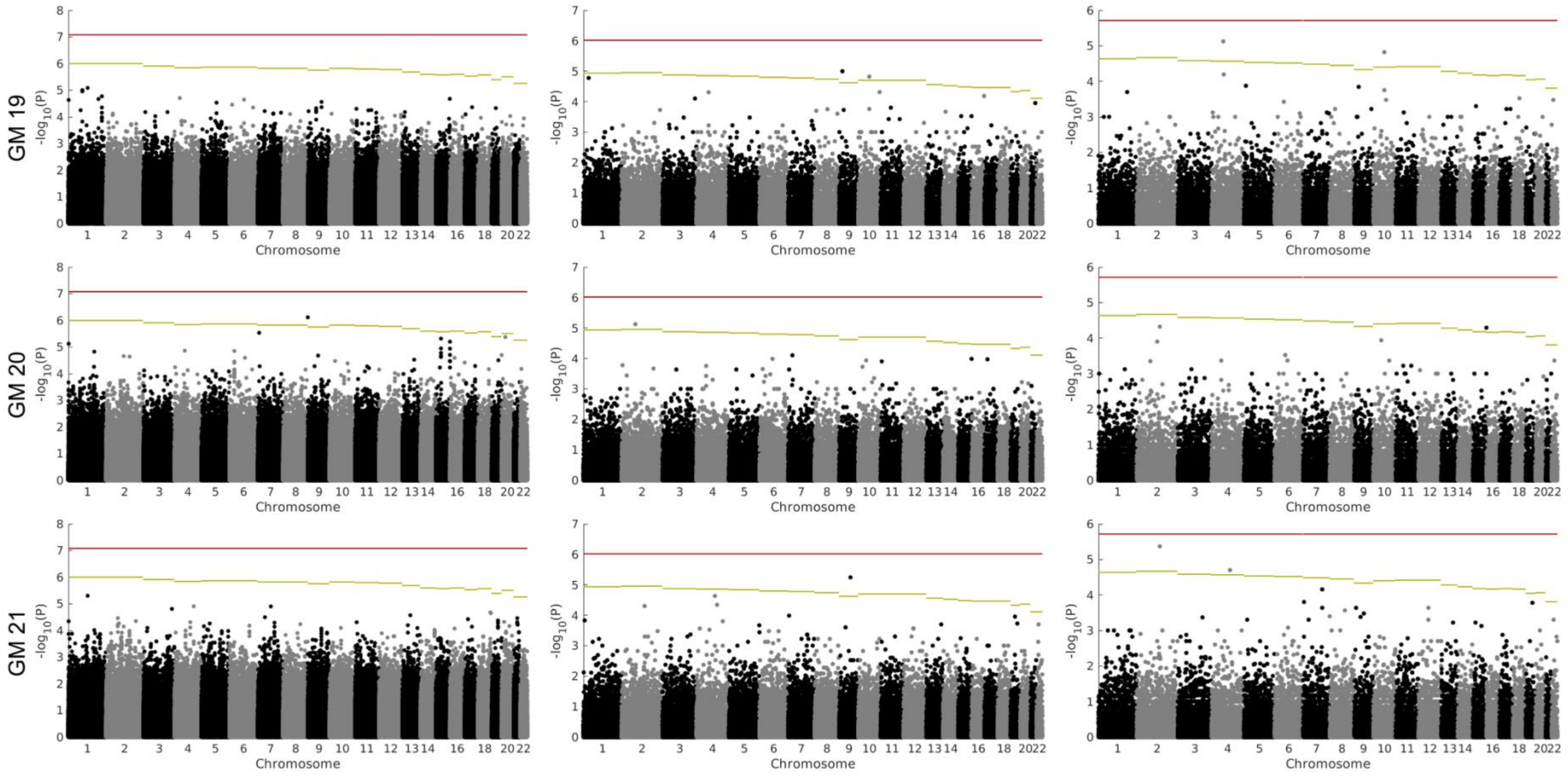
GM 19
GM 20
GM 21
-log10(P)
Chromosome

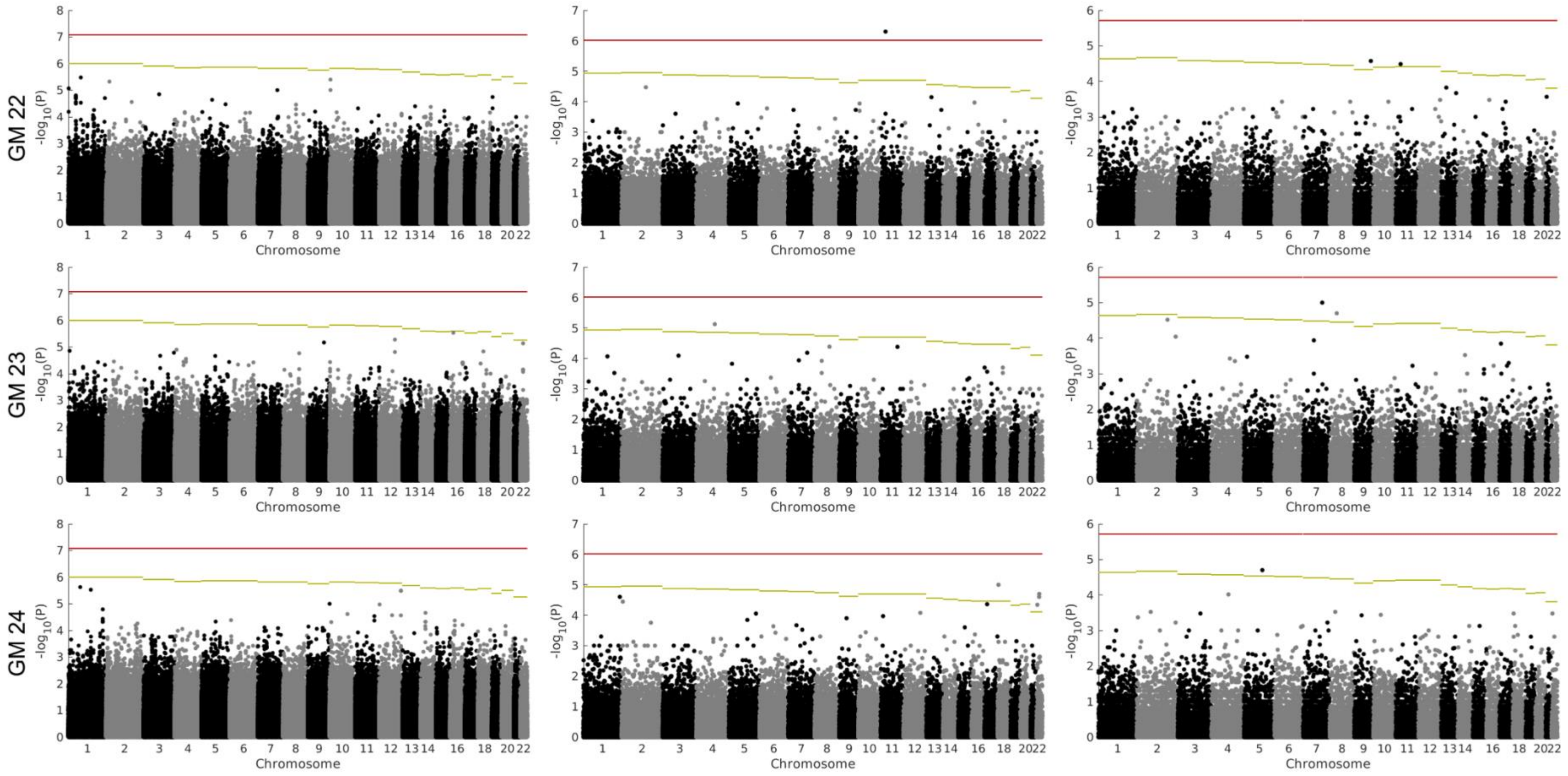

GM 22
GM 23
GM 24
-log10(P)
Chromosome

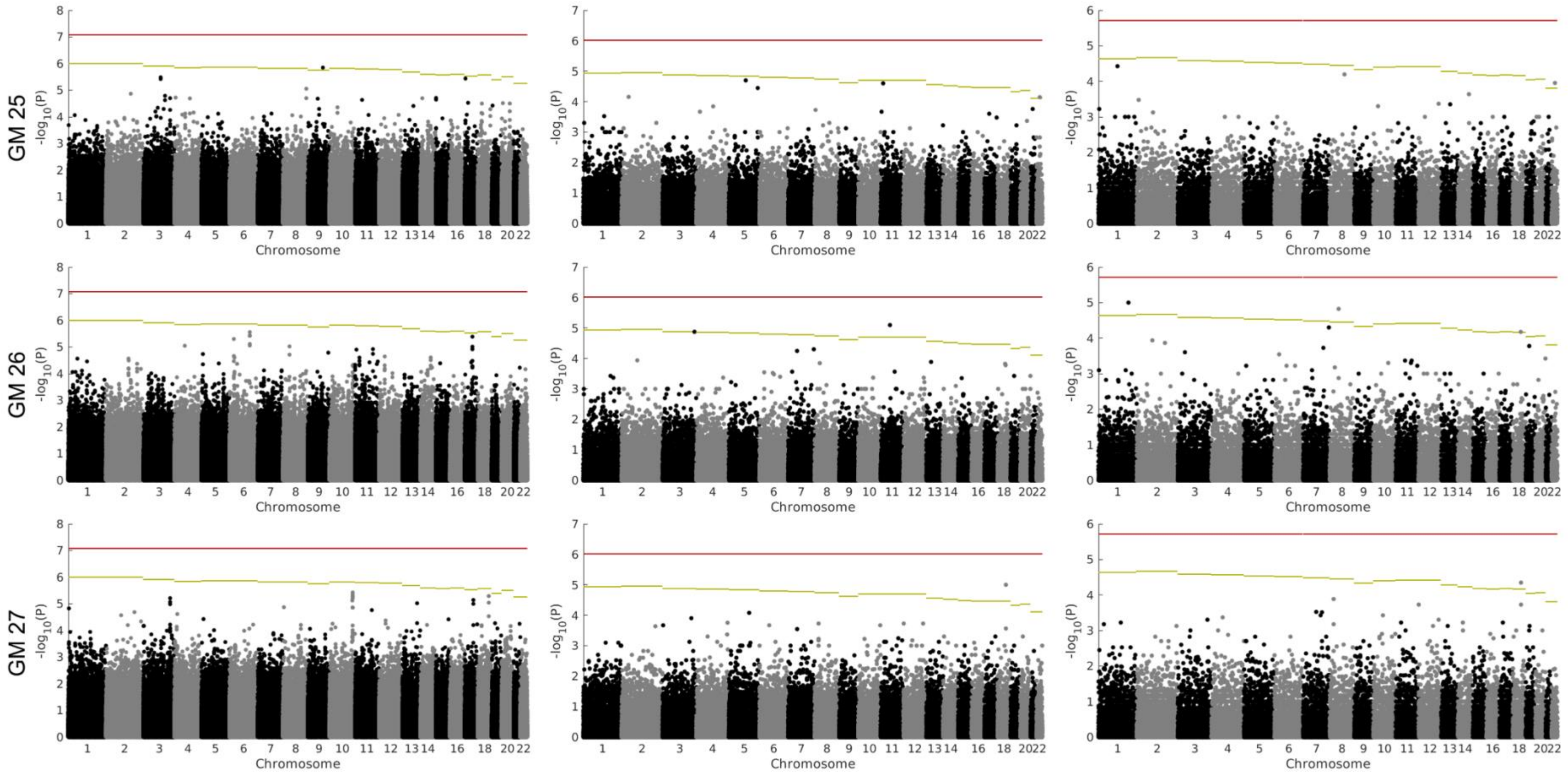
GM 25
GM 26
GM 27
-log10(P)
Chromosome
1 2 3 4 5 6 7 8 9 10 11 12 13 14 16 18 20 22

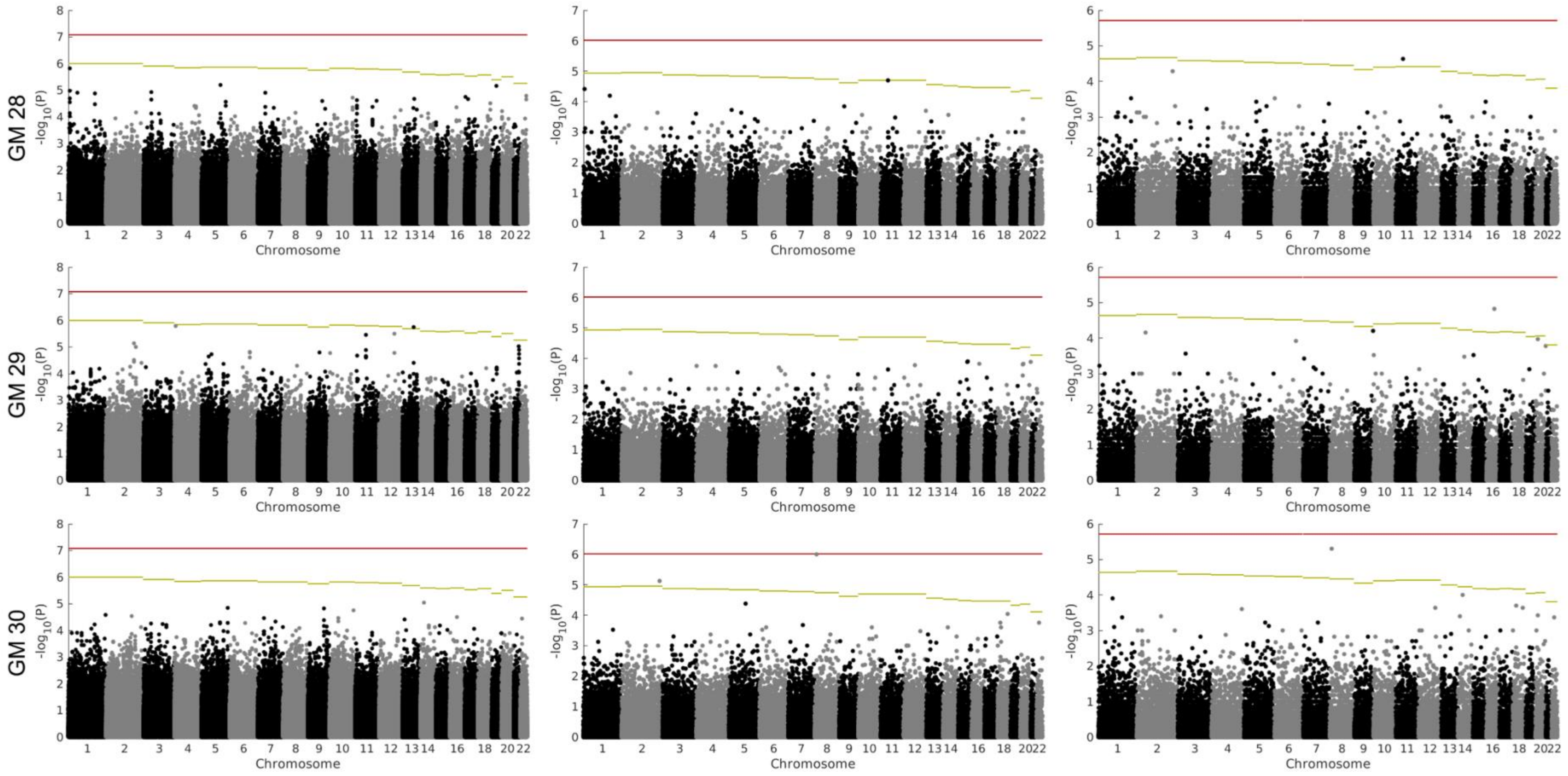

GM 28
GM 29
GM 30
-log10(P)
Chromosome

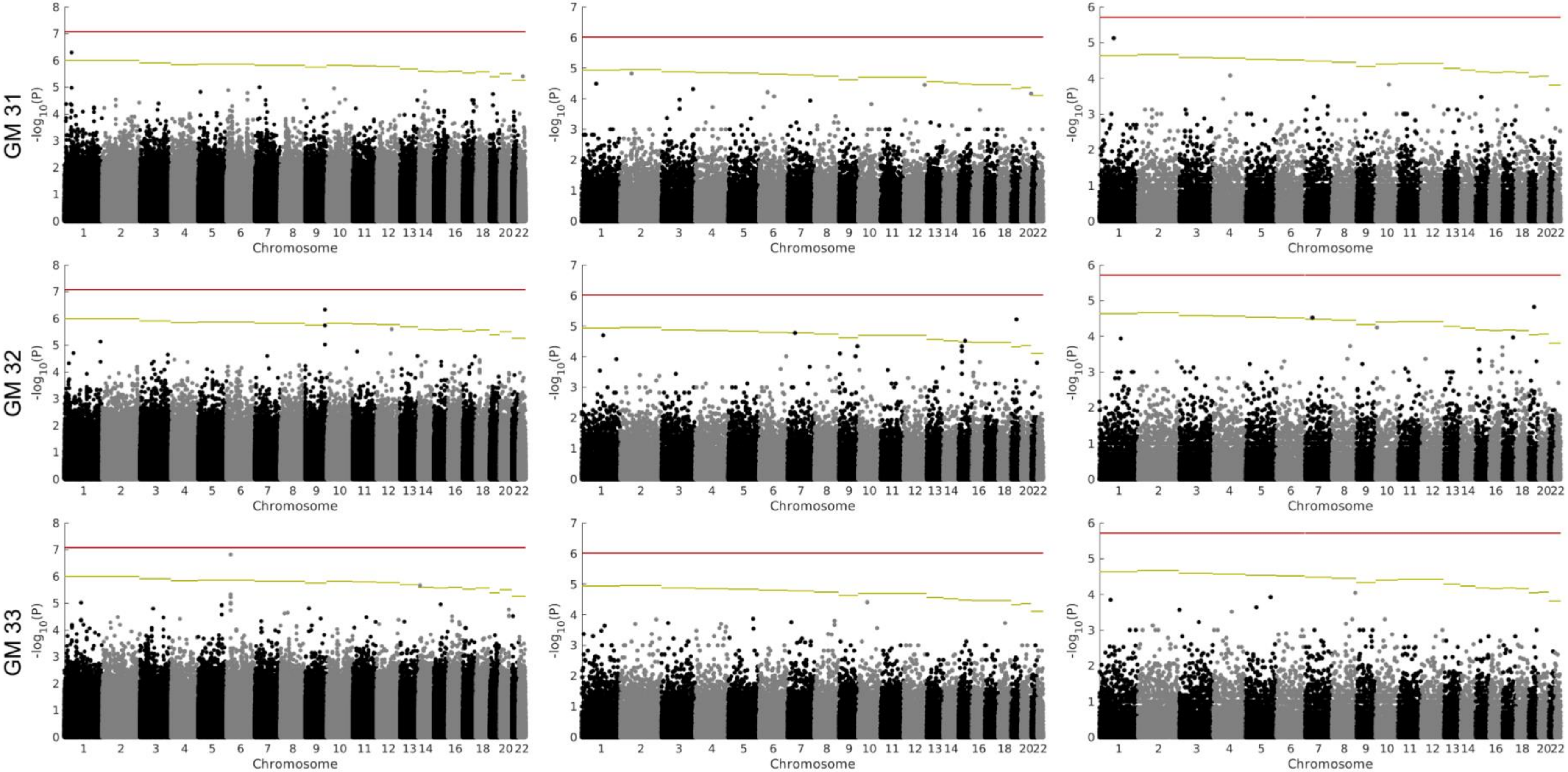
GM 31
GM 32
GM 33
-log10(P)
Chromosome
1 2 3 4 5 6 7 8 9 10 11 12 13 14 16 18 20 22

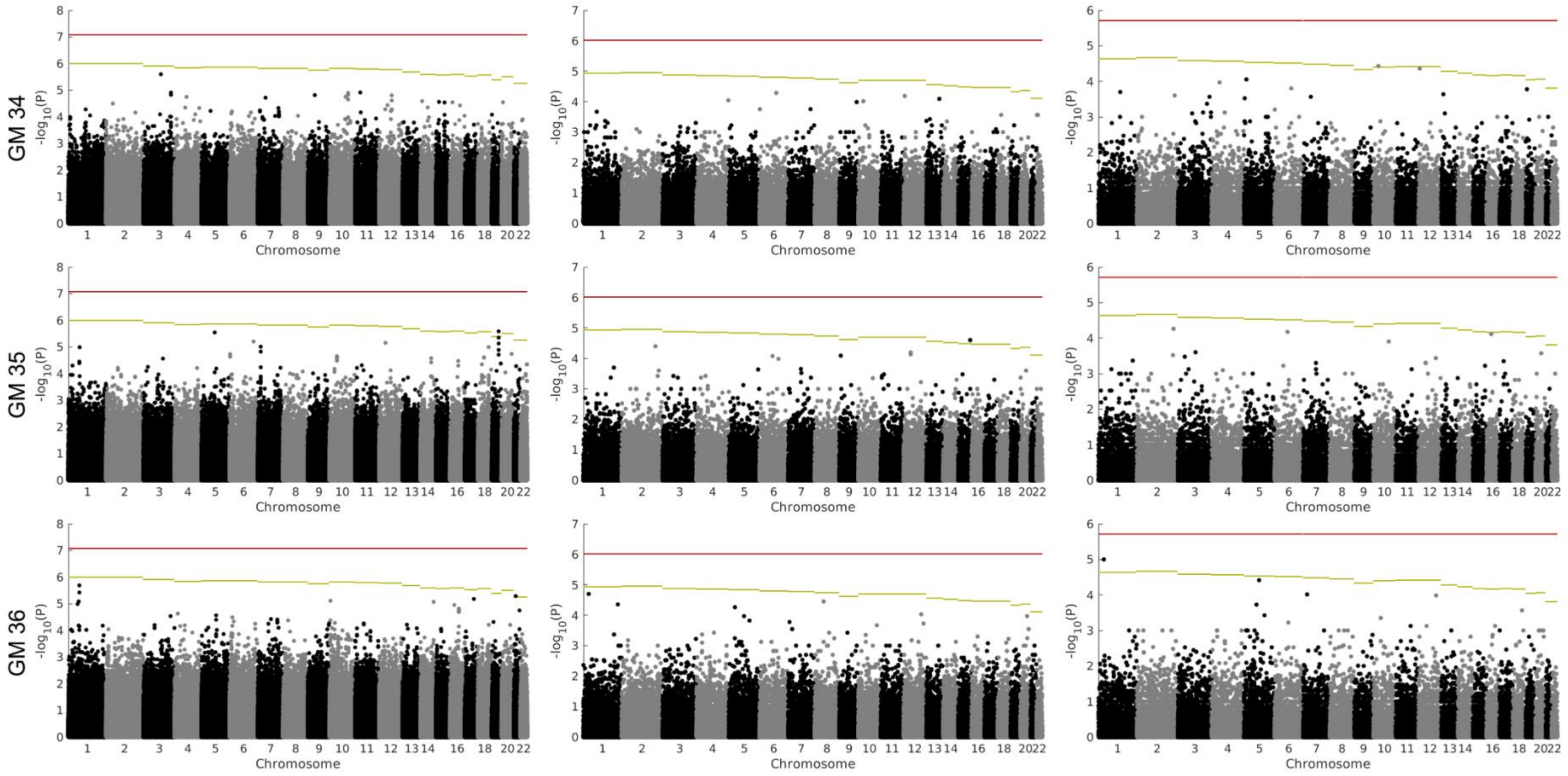
GM 34
GM 35
GM 36
-log10(P)
Chromosome

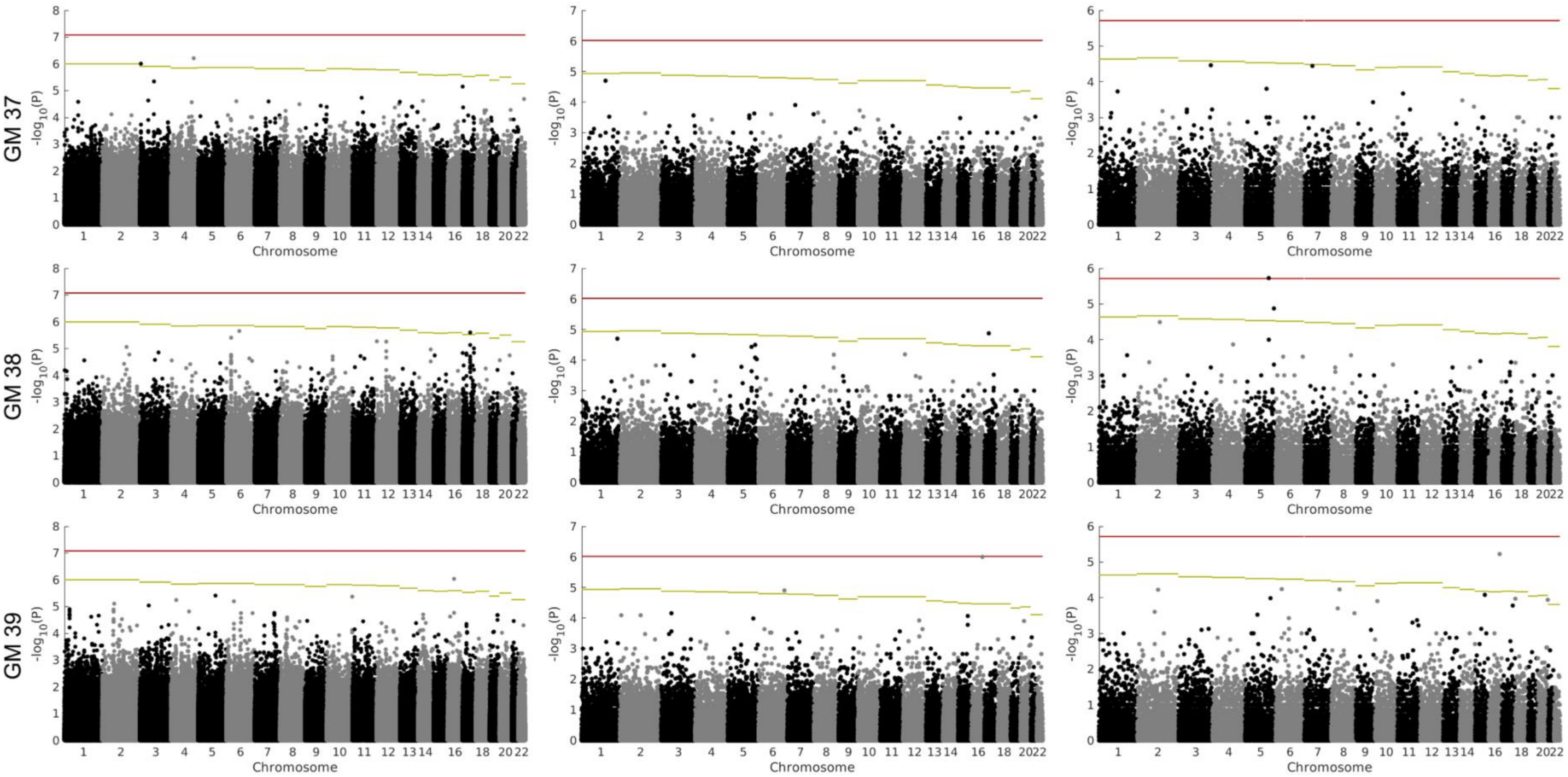
GM 37
GM 38
GM 39
-log10(P)
Chromosome

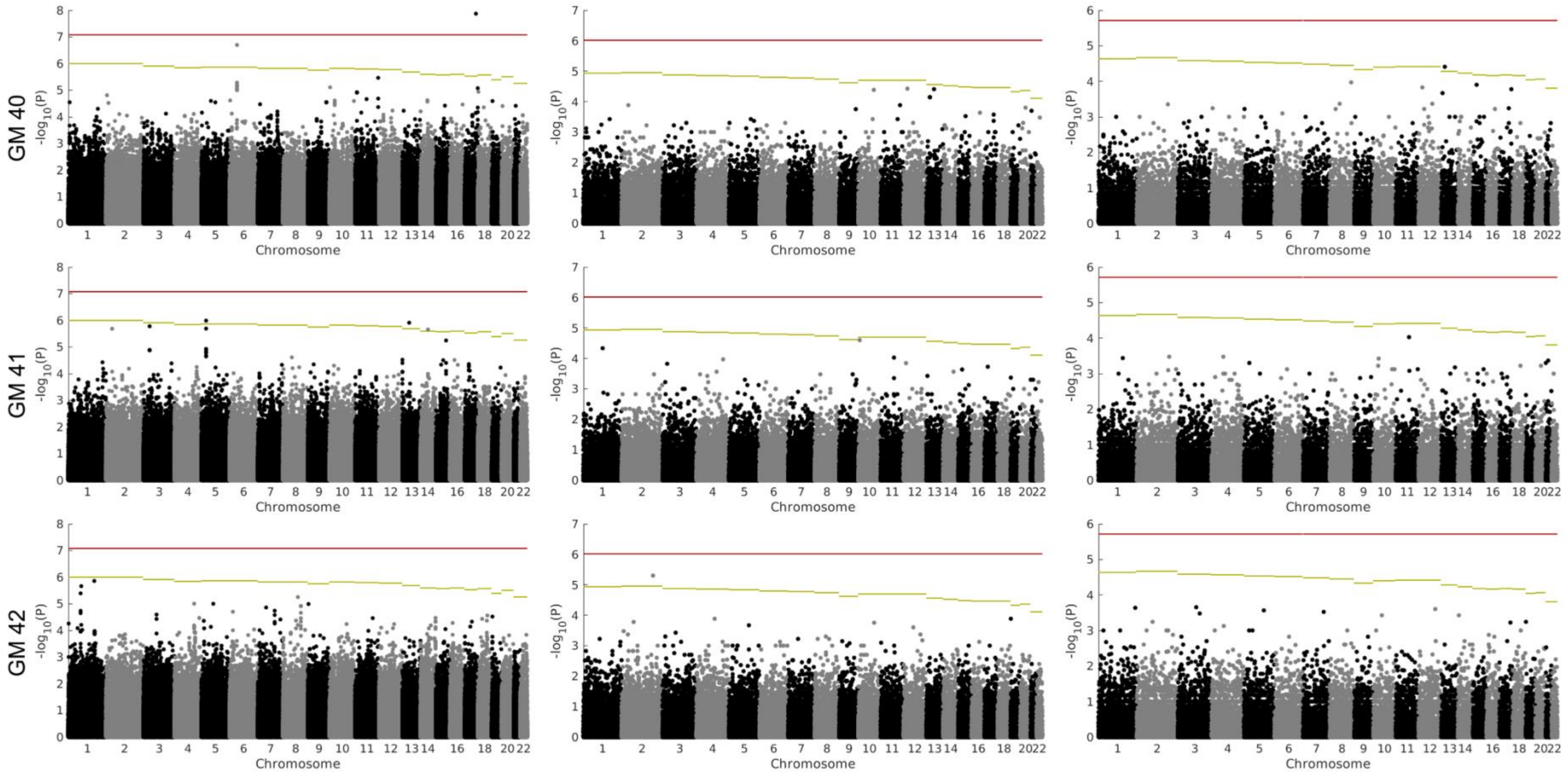
GM 40
GM 41
GM 42
-log10(P)
Chromosome
1 2 3 4 5 6 7 8 9 10 11 12 13 14 16 18 20 22

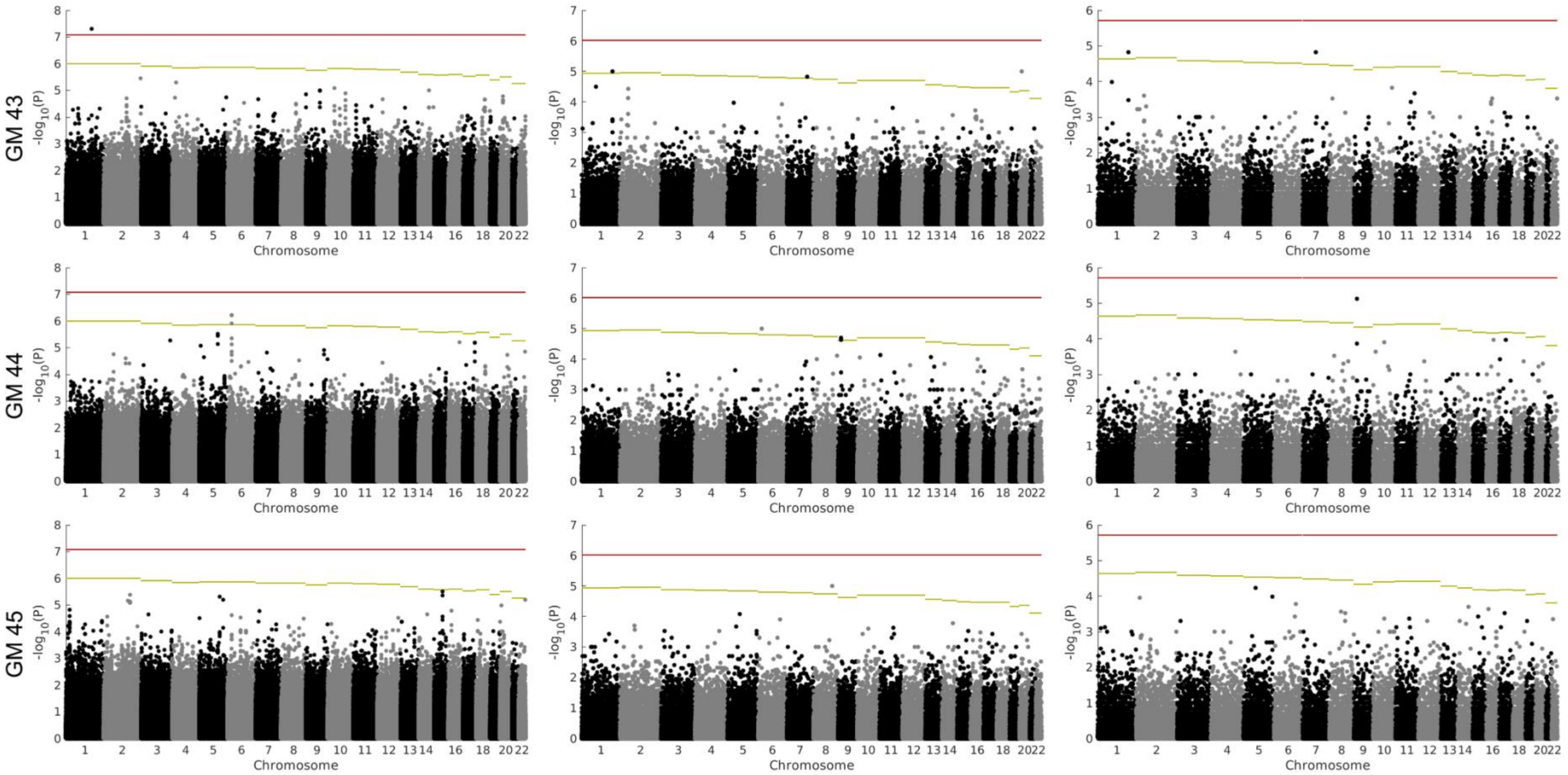
GM 43
GM 44
GM 45
-log10(P)
Chromosome

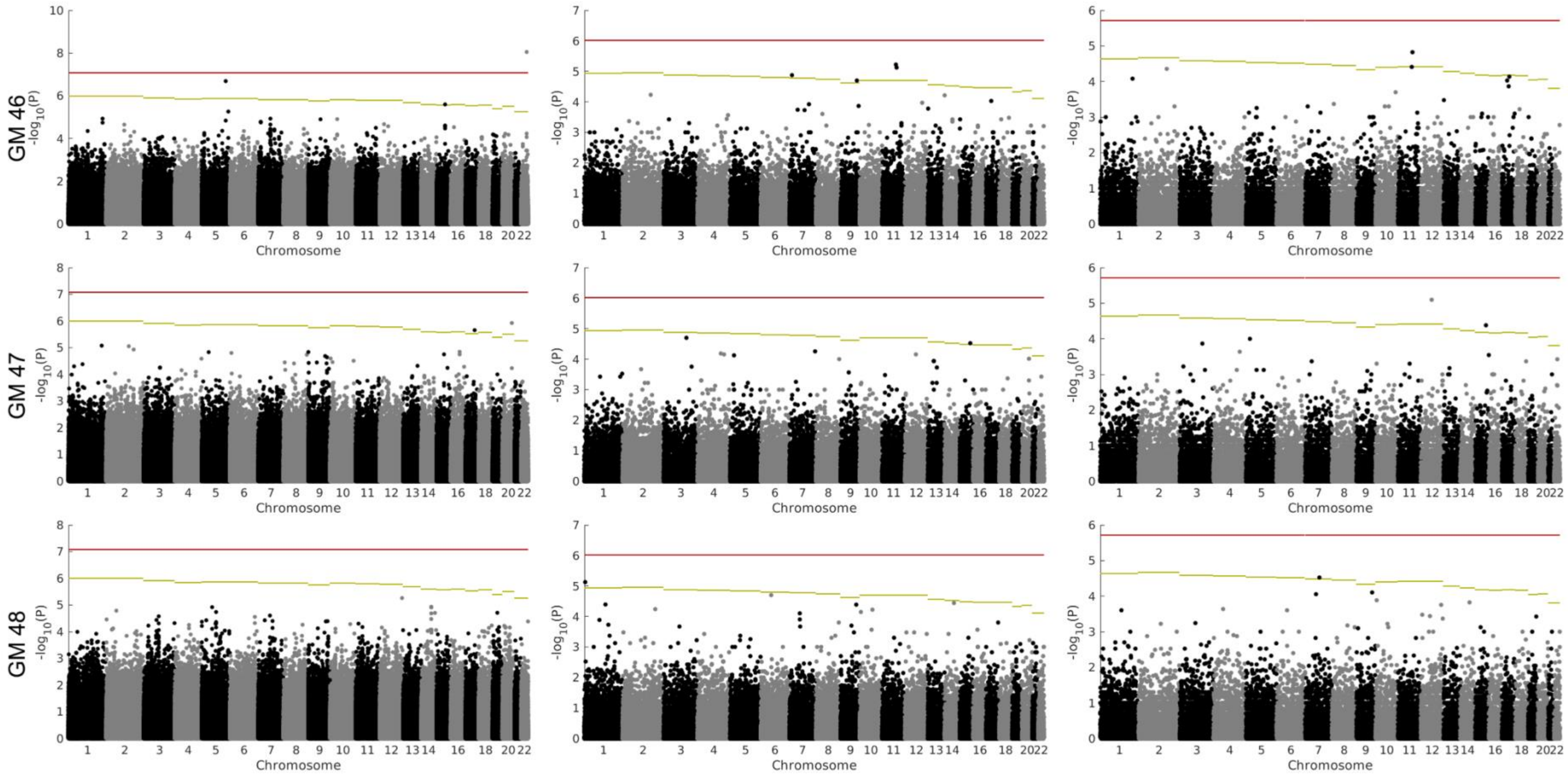
GM 46
GM 47
GM 48
-log10(P)
Chromosome

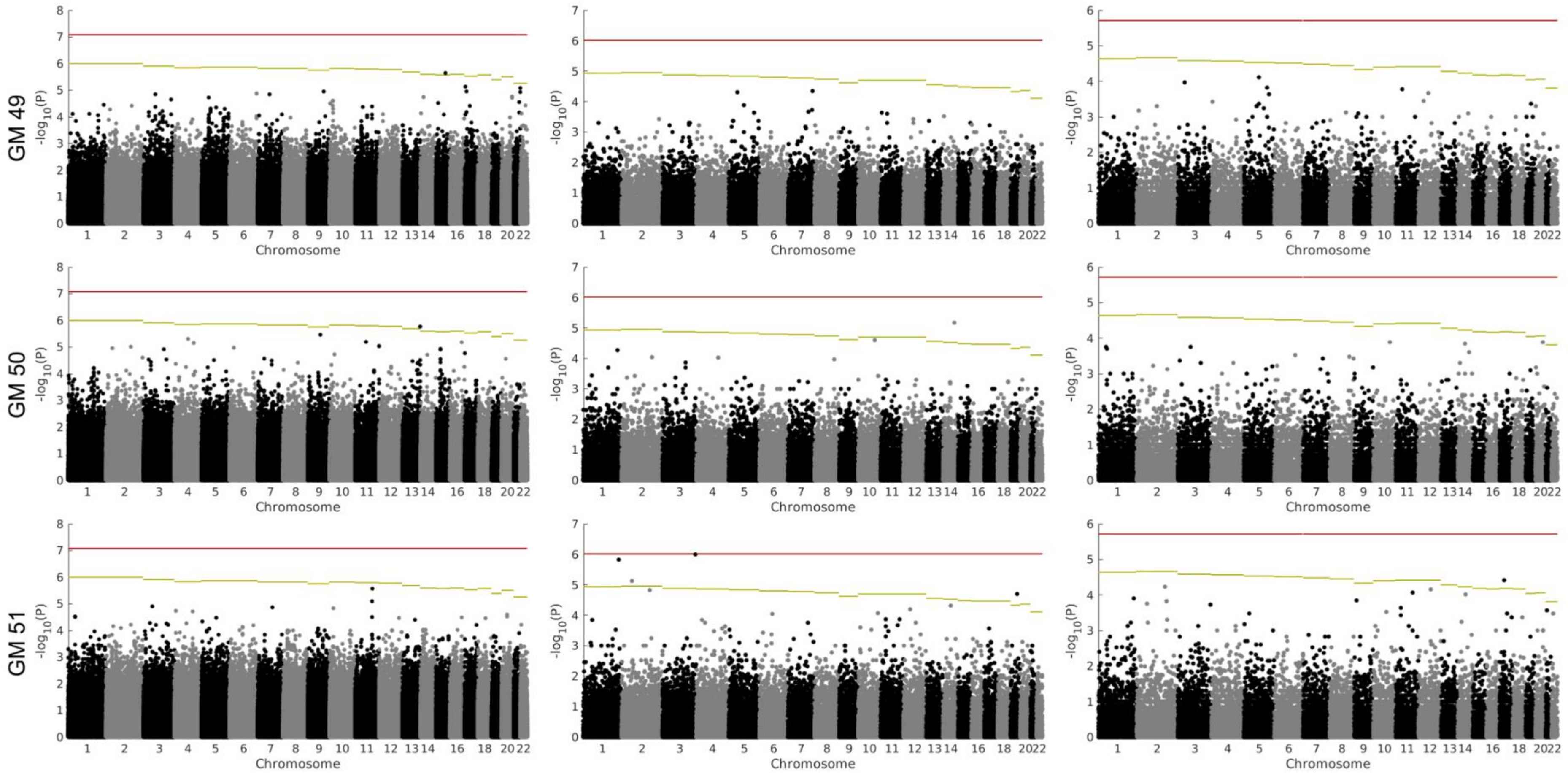
GM 49
GM 50
GM 51
-log10(P)
Chromosome

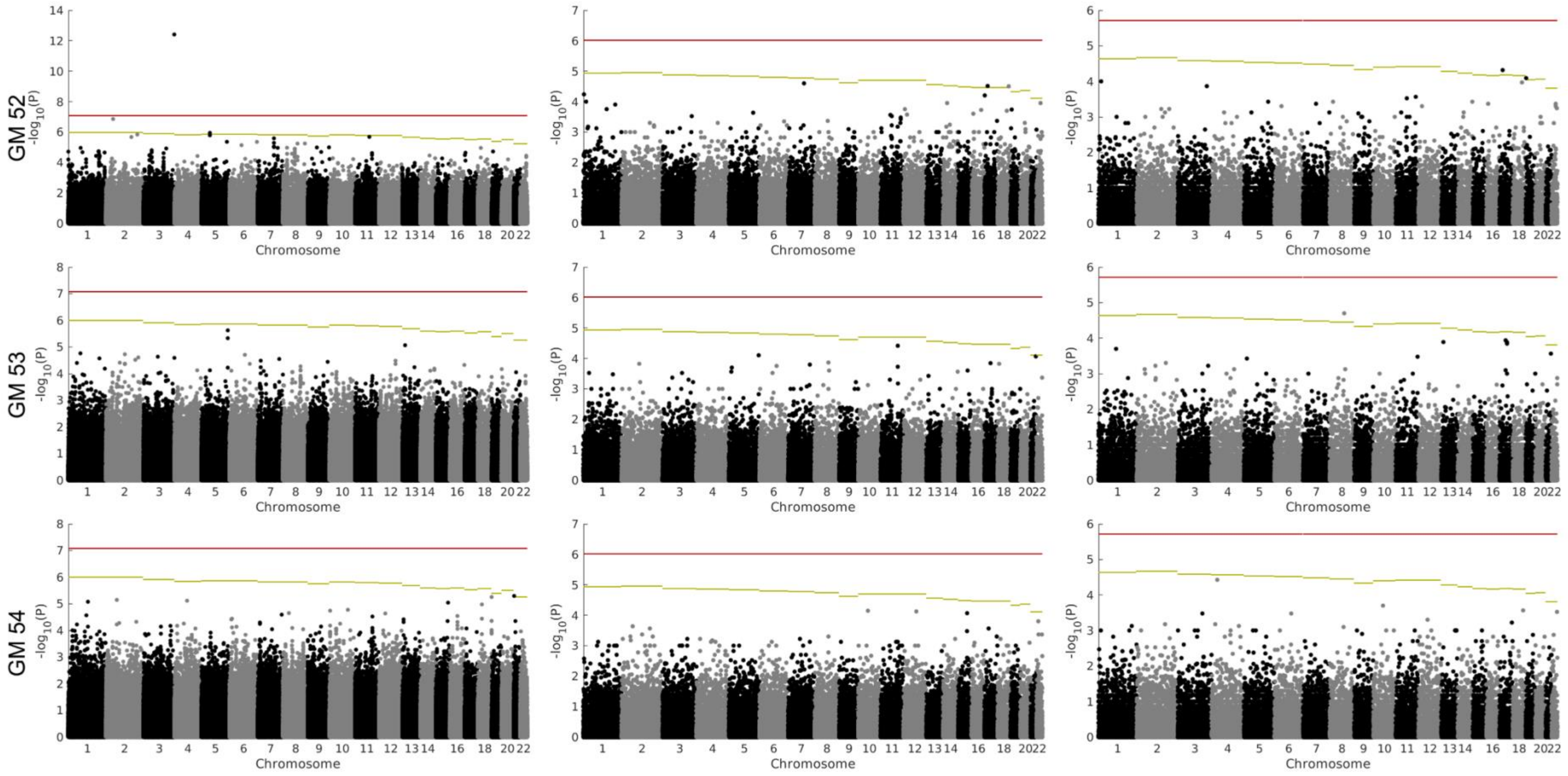
GM 52
GM 53
GM 54
-log10(P)
Chromosome

GM 55

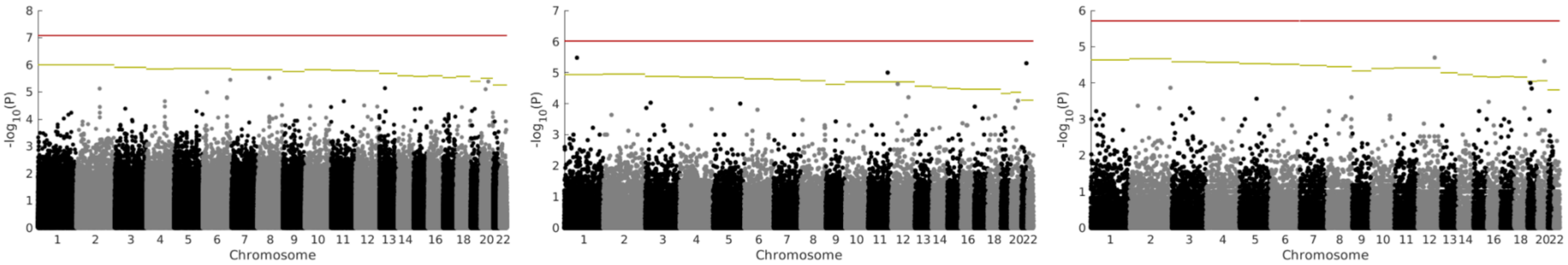

**Figure S2** GWAS and REL results of white matter tract fractional anisotropy (PC1, PC2 and 10 tracts). Left: GWAS results (Wald p-value), middle & right: REL results of 50kb & 100kb genomic windows (Permutation p-value). Red horizontal bar: genome-wide Bonferroni correction threshold. Yellow horizontal bar: chromosome-wide Bonferroni correction threshold.

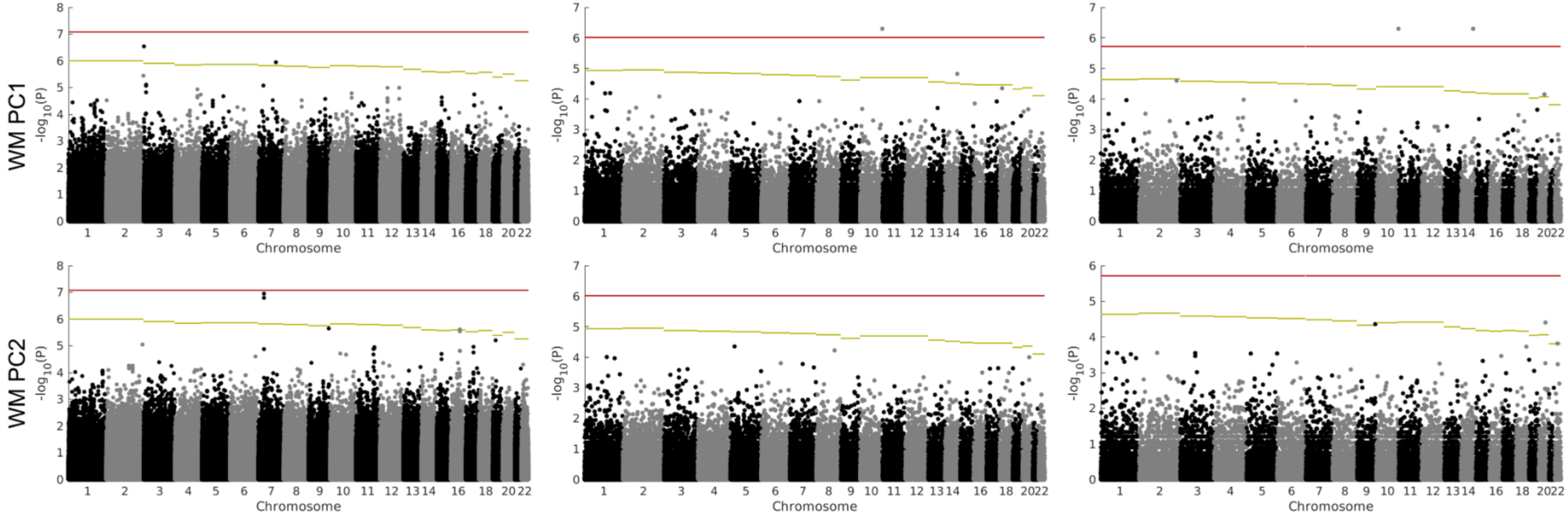

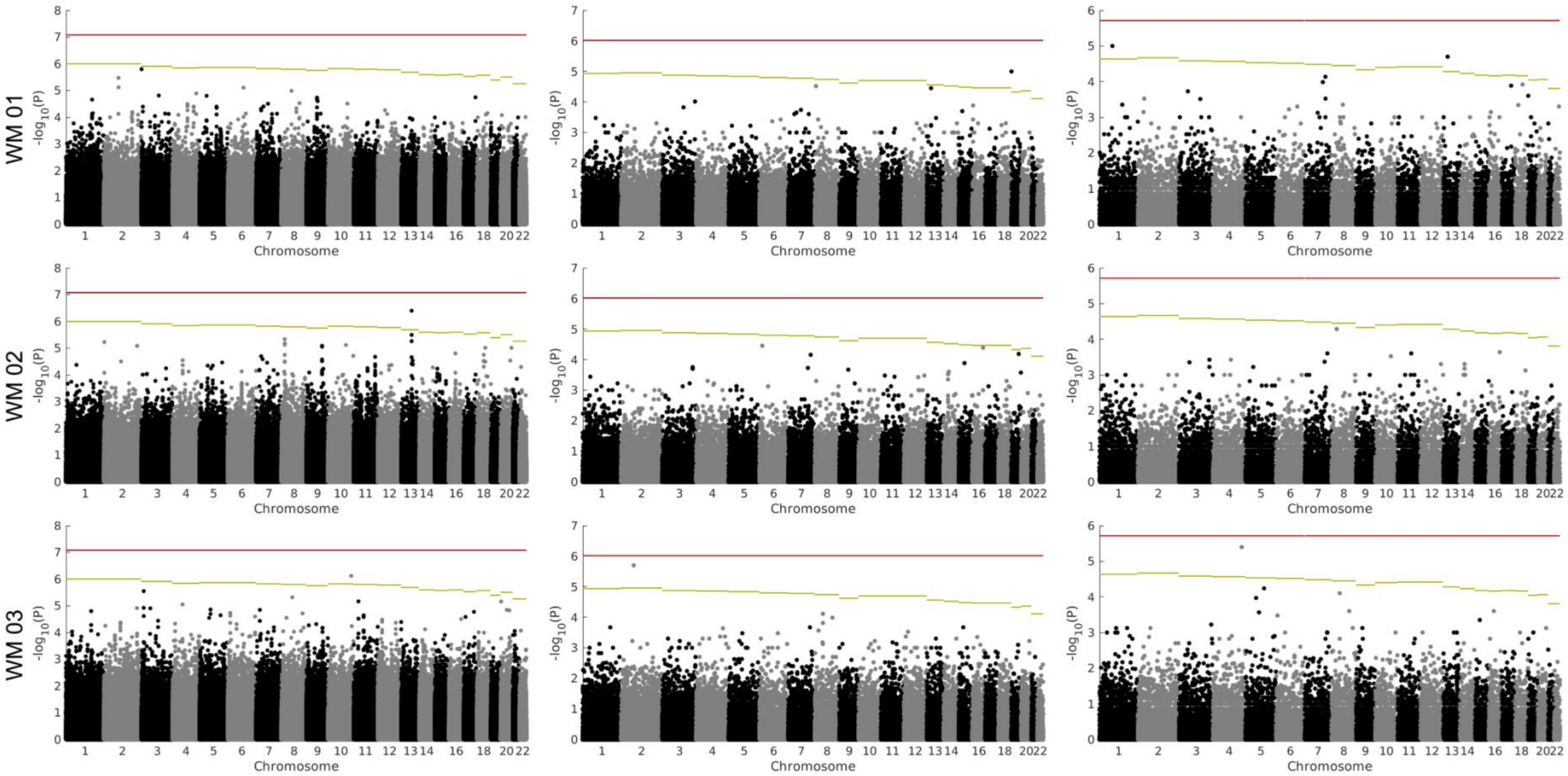
WM 01
WM 02
WM 03
-log10(P)
Chromosome

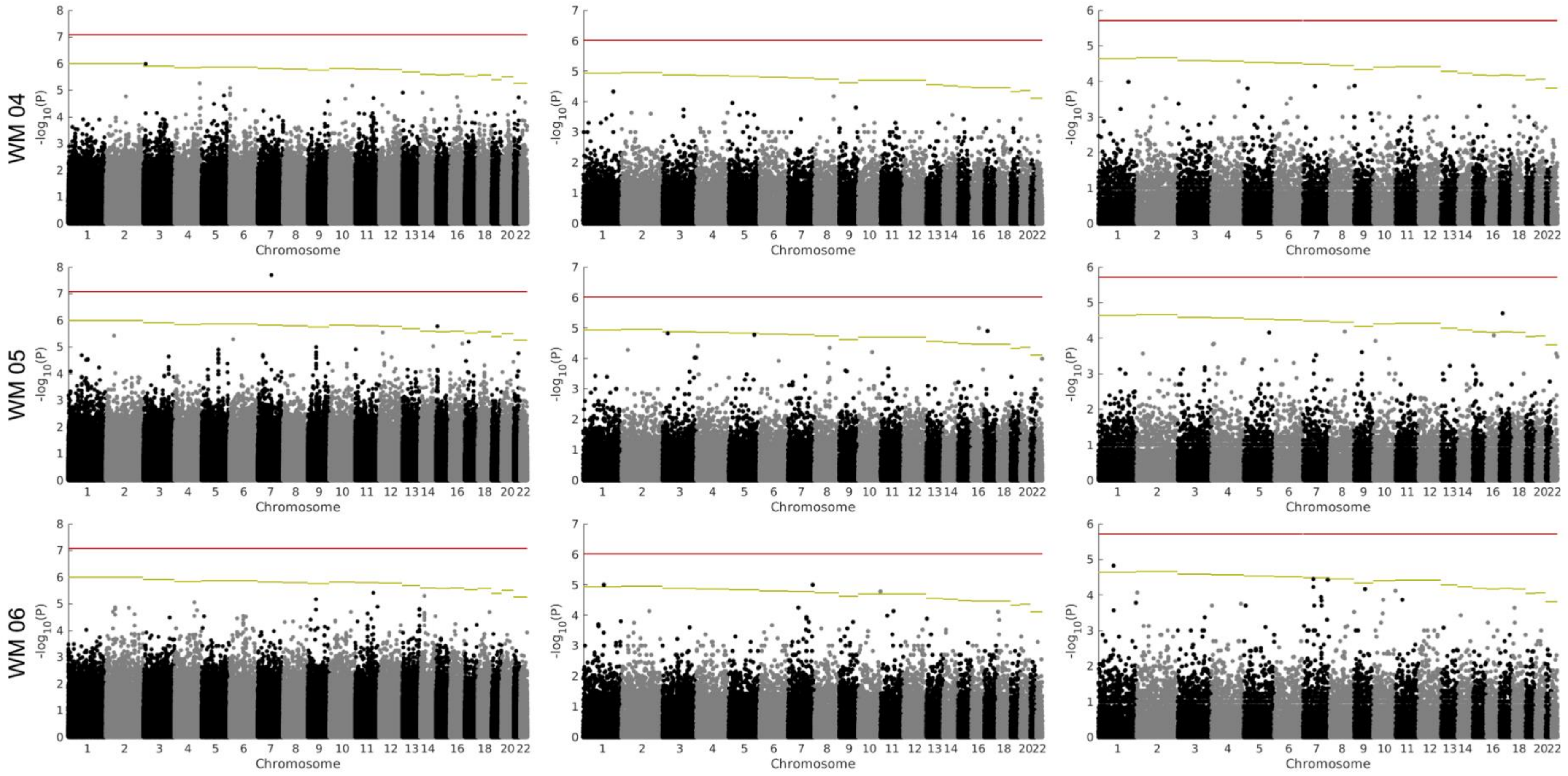
WM 04
WM 05
WM 06
-log10(P)
Chromosome

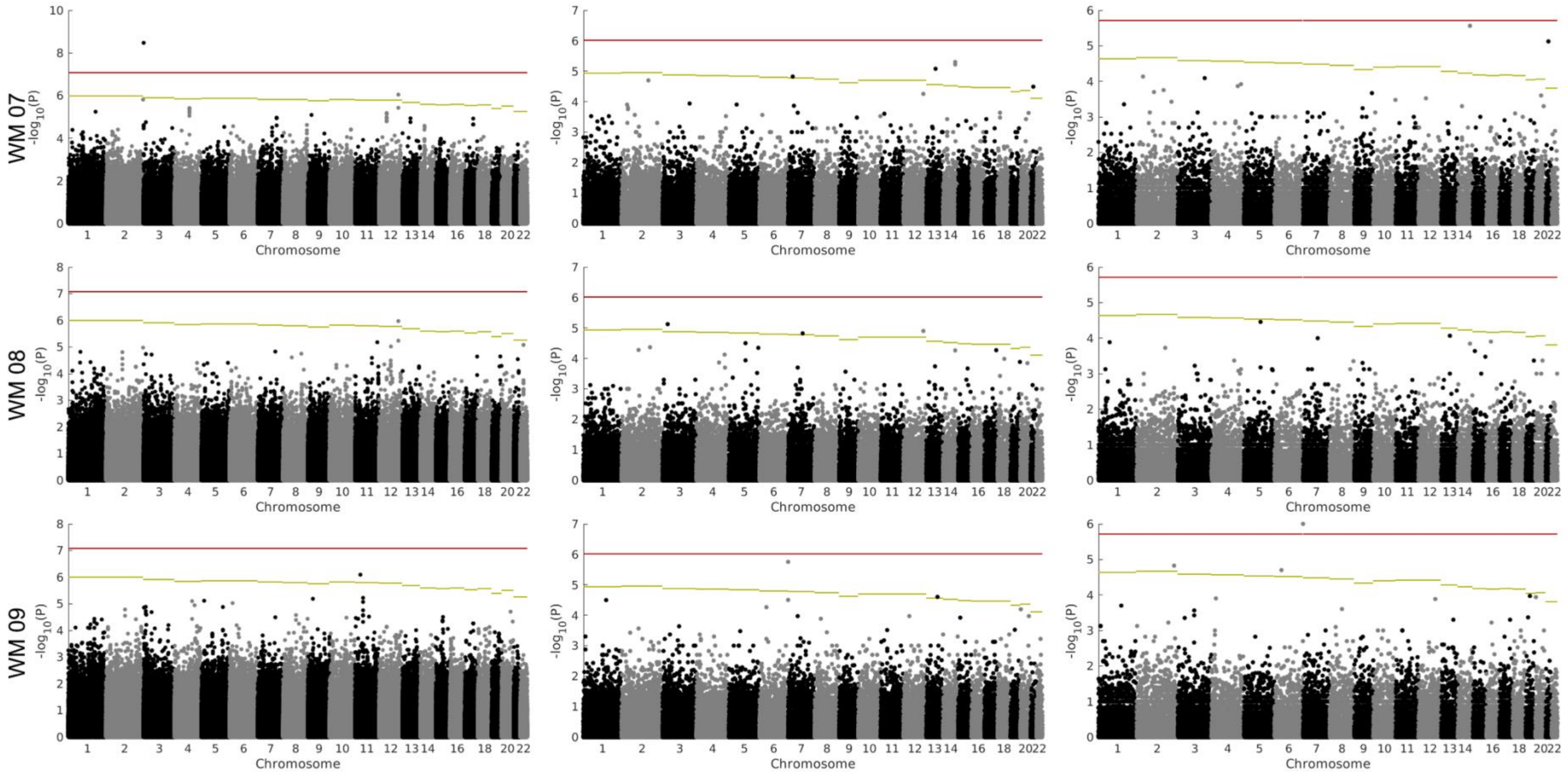

WM 07
WM 08
WM 09
-log10(P)
Chromosome

WM 10

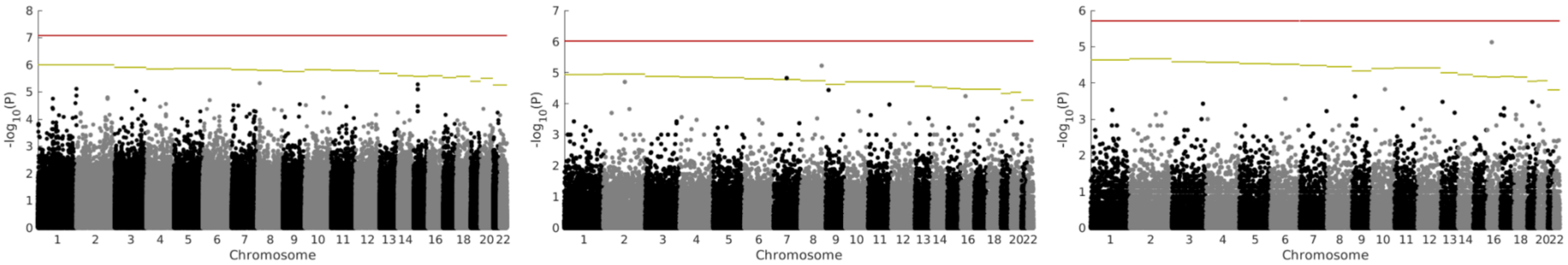
