## Supplemental Tables 1-4 for "Genetic signatures of human brain structure: A comparison between GWAS and relatedness-based regression"

**Table S1** Harvard-Oxford cortical and subcortical structural atlases

| Index | Abbreviation | Full description | Brodmann Area |
|---|---|---|---|
| 1 | FP | Frontal Pole | 10 |
| 2 | Ins | Insular Cortex | N/A |
| 3 | SFG | Superior Frontal Gyrus | 6 |
| 4 | MFG | Middle Frontal Gyrus | 9 |
| 5 | IFGpt | Inferior Frontal Gyrus, pars triangularis | 45 |
| 6 | IFGpo | Inferior Frontal Gyrus, pars opercularis | 44 |
| 7 | PrG | Precentral Gyrus | 6 |
| 8 | TP | Temporal Pole | 38 |
| 9 | STGa | Superior Temporal Gyrus, anterior division | 22 |
| 10 | STGp | Superior Temporal Gyrus, posterior division | 22 |
| 11 | MTGa | Middle Temporal Gyrus, anterior division | 21 |
| 12 | MTGp | Middle Temporal Gyrus, posterior division | 21 |
| 13 | MTGtp | Middle Temporal Gyrus, temporooccipital part | 21/22 |
| 14 | ITGa | Inferior Temporal Gyrus, anterior division | 20 |
| 15 | ITGp | Inferior Temporal Gyrus, posterior division | 20 |
| 16 | ITGtp | Inferior Temporal Gyrus, temporooccipital part | 20/22 |
| 17 | PoG | Postcentral Gyrus | 3/4 |
| 18 | SPL | Superior Parietal Lobule | 40 |
| 19 | SmGa | Supramarginal Gyrus, anterior division | 40 |
| 20 | SmGp | Supramarginal Gyrus, posterior division | 40 |
| 21 | AG | Angular Gyrus | 40 |
| 22 | LOCs | Lateral Occipital Cortex, superior division | 19 |
| 23 | LOCi | Lateral Occipital Cortex, inferior division | 19 |
| 24 | IcC | Intracalcarine Cortex | 23 |
| 25 | FMC | Frontal Medial Cortex | 11 |
| 26 | SMC | Juxtapositional Lobule Cortex (formerly Supplementary Motor Cortex) | 6 |
| 27 | ScC | Subcallosal Cortex | 25 |
| 28 | PcG | Paracingulate Gyrus | 32 |
| 29 | CGa | Cingulate Gyrus, anterior division | 24/33 |
| 30 | CGp | Cingulate Gyrus, posterior division | 23/31 |
| 31 | PcC | Precuneous Cortex | 7 |
| 32 | CC | Cuneal Cortex | 18 |
| 33 | FOC | Frontal Orbital Cortex | 47 |
| 34 | PhGa | Parahippocampal Gyrus, anterior division | 35 |
| 35 | PaGp | Parahippocampal Gyrus, posterior division | 36 |
| 36 | LG | Lingual Gyrus | 18 |
| 37 | TFCa | Temporal Fusiform Cortex, anterior division | 20 |
| 38 | TFCp | Temporal Fusiform Cortex, posterior division | 36 |
| 39 | TOF | Temporal Occipital Fusiform Cortex | 37 |
| 40 | OFG | Occipital Fusiform Gyrus | 19 |
| 41 | FOpC | Frontal Operculum Cortex | 13 |
| 42 | COpC | Central Opercular Cortex | 6/13 |
| 43 | POpC | Parietal Operculum Cortex | 13 |
| 44 | PP | Planum Polare | 22 |
| 45 | H1/H2 | Heschl's Gyrus (includes H1 and H2) | 13 |
| 46 | PT | Planum Temporale | 41 |
| 47 | SccC | Supracalcarine Cortex | 18 |
| 48 | OcP | Occipital Pole | 18 |
| 49 | MDN | Medial Dorsal Neucleus | N/A |
| 50 | CauB | Caudate Body | N/A |
| 51 | Puta | Putamen | N/A |
| 52 | LGP | Lateral Globus Pallidus | N/A |
| 53 | Hipp | Hippocampus | 28 |
| 54 | Amy | Amygdala | N/A |
| 55 | CauH | Caudate Head | N/A |

**Table S2** JHU DTI-based white-matter atlases

| Index | Full description |
|---|---|
| 1 | Anterior Thalamic Radiation |
| 2 | Corticospinal Tract |
| 3 | Cingulum (cingulate gyrus) |
| 4 | Cingulum (hippocampus) |
| 5 | Forceps Major |
| 6 | Forceps Minor |
| 7 | Inferior Fronto-occipital Fasciculus |
| 8 | Inferiorongitudinal Fasciculus |
| 9 | Superiorongitudinal Fasciculus |
| 10 | Uncinate Fasciculus |
| 11 | Superiorongitudinal Fasciculus (temporal part) |

**Table S3** Loci associated with brain stratural measures in GWAS

| | phenotype | chr | SNP | position (KB) | Alleles | P_wald | Related gene |
|---|---|---|---|---|---|---|---|
| GM_05 | Inferior Frontal Gyrus, pars triangularis | 15 | rs6494798 | 69790.976 | C>G,T | 2.89E-08 | N/A |
| GM_05 | Inferior Frontal Gyrus, pars triangularis | 15 | rs11634551 | 69808.89 | C>T | 6.99E-09 | N/A |
| GM_05 | Inferior Frontal Gyrus, pars triangularis | 15 | rs6494801 | 69822.411 | G>A,T | 6.99E-09 | N/A |
| GM_17 | Postcentral Gyrus | 19 | rs10424191 | 31057.349 | A>G | 4.10E-08 | ZNF536 |
| GM_40 | Occipital Fusiform Gyrus | 17 | rs62621376 | 72233.559 | G>A | 1.33E-08 | TTYH2 |
| GM_43 | Parietal Operculum Cortex | 1 | rs872376 | 179794.602 | A>G | 4.94E-08 | LINC02818 |
| GM_46 | Planum Temporale | 22 | rs5765545 | 46022.046 | C>T | 8.87E-09 | N/A |
| GM_52 | Lateral Globus Pallidus | 3 | rs36016914 | 196050.862 | A>G | 3.88E-13 | TM4SF19, TM4SF19-TCTEX1D2, TM4SF19-AS1 |
| WM_05 | Forceps Major | 7 | rs143118835 | 80141.152 | C>T | 1.99E-08 | GNAT3, LOC107986812 |
| WM_07 | Inferior Fronto-occipital Fasciculus | 3 | rs9882746 | 595.87 | G>A | 3.36E-09 | N/A |

SNP position is based on the reference assembly sequence GRCh37.

**Table S4** Genomic windows associated with brain stratural measures in REL

| | phenotype | chr | window width (KB) | window onset (KB) | window offset (KB) | P_perm | number of SNPs | SNPs | Related gene |
|---|---|---|---|---|---|---|---|---|---|
| GM PC1 | global measure | 17 | 50 | 37160.446 | 37207.734 | 5.00E-07 | 6 | rs1014263, rs11868358, rs9914342, rs8071973, rs12600916, rs11868463 | LRRC37A11P, LOC105371767 |
| GM PC1 | global measure | 17 | 100 | 37152.369 | 37207.734 | 5.00E-07 | 7 | rs8070153, rs1014263, rs11868358, rs9914342, rs8071973, rs12600916, rs11868463 | LRRC37A11P, LOC105371767 |
| GM_07 | Precentral Gyrus | 16 | 50 | 20952.098 | 20982.832 | 5.00E-07 | 16 | rs11649212, rs111171, rs2301620, rs428863, rs147732992, rs12051478, rs34771199, rs34051490, rs370719, rs111539520, rs3743696, rs3743697, rs61735055, rs33928718, rs3103824, rs7189852 | DNAH3 |
| GM_07 | Precentral Gyrus | 16 | 100 | 20909.404 | 20982.832 | 5.00E-07 | 24 | rs12325560, rs3815019, rs17692345, rs2269766, rs17692528, rs11864905, rs9925327, rs4783513, rs11649212, rs111171, rs2301620, rs428863, rs147732992, rs12051478, rs34771199, rs34051490, rs370719, rs111539520, rs3743696, rs3743697, rs61735055, rs33928718, rs3103824, rs7189852 | DNAH3, LYRM1, DCUN1D3 |
| GM_13 | Middle Temporal Gyrus, temporooccipital part | 14 | 50 | 33881.216 | 33926.191 | 5.00E-07 | 30 | rs17101040, rs10148313, rs8018275, rs17101062, rs1440198, rs11622370, rs10134290, rs10129955, rs11621942, rs17101124, rs967440, rs12147692, rs12147072, rs8018440, rs11156790, rs10141664, rs10133550, rs4981191, rs12897109, rs17101176, rs10134359, rs927327, rs4981192, rs8014355, rs8006708, rs1440197, rs8012273, rs8003435, rs1030976, rs1030974 | NPAS3 |
| GM_13 | Middle Temporal Gyrus, temporooccipital part | 14 | 100 | 33829.28 | 33926.191 | 1.00E-06 | 62 | rs2274512, rs2274511, rs1315131, rs1315130, rs10483439, rs1948724, rs1315128, rs17100828, rs11156787, rs17100845, rs17491398, rs10148795, rs1315148, rs1110868, rs11851892, rs10144759, rs1887496, rs10143306, rs1604190, rs1303470, rs10483440, rs1595836, rs1117363, rs4982083, rs7161650, rs17101006, rs1160163, rs3742932, rs17101028, rs10047941, rs1371707, rs4982084, rs17101040, rs10148313, rs8018275, rs17101062, rs1440198, rs11622370, rs10134290, rs10129955, rs11621942, rs17101124, rs967440, rs12147692, rs12147072, rs8018440, rs11156790, rs10141664, rs10133550, rs4981191, rs12897109, rs17101176, rs10134359, rs927327, rs4981192, rs8014355, rs8006708, rs1440197, rs8012273, rs8003435, rs1030976, rs1030974 | NPAS3 |
| GM_17 | Postcentral Gyrus | 8 | 100 | 61866.711 | 61964.013 | 5.00E-07 | 36 | rs11781582, rs12335258, rs4412366, rs13280897, rs882229, rs12678105, rs6992440, rs10113311, rs7842384, rs6471907, rs2280813, rs9692725, rs16926675, rs10957177, rs7841986, rs3864678, rs6989470, rs3864664, rs3963188, rs10504318, rs16926700, rs6995339, rs12542604, rs6986434, rs7004557, rs7814400, rs1962862, rs1871217, rs1465667, rs11986500, rs6471908, rs7827230, rs2882217, rs10097565, rs4738858, rs16926764 | N/A |
| GM_17 | Postcentral Gyrus | 16 | 100 | 20909.404 | 20982.832 | 1.67E-06 | 24 | rs12325560, rs3815019, rs17692345, rs2269766, rs17692528, rs11864905, rs9925327, rs4783513, rs11649212, rs111171, rs2301620, rs428863, rs147732992, rs12051478, rs34771199, rs34051490, rs370719, rs111539520, rs3743696, rs3743697, rs61735055, rs33928718, rs3103824, rs7189852 | DNAH3, LYRM1, DCUN1D3 |
| GM_22 | Lateral Occipital Cortex, superior division | 11 | 50 | 27946.346 | 27990.119 | 5.00E-07 | 7 | rs10835237, rs2726832, rs1404950, rs2726829, rs1464896, rs12799489, rs10501093 | N/A |
| GM_38 | Temporal Fusiform Cortex, posterior division | 5 | 100 | 140653.258 | 140735.215 | 1.88E-06 | 33 | rs10044936, rs618578, rs619268, rs2879202, rs1422411, rs1076361, rs11740372, rs6580184, rs13187825, rs10491308, rs10491309, rs7735993, rs11952797, rs6898211, rs17097145, rs11547633, rs10875595, rs13359820, rs76571170, rs17097187, rs11575968, rs10491311, rs6861047, rs61749056, rs66823521, rs3806837, rs11575948, rs72790006, rs17097224, rs62378414, rs77250251, rs10214288, rs11575949 | PCDHGA4, PCDHGA3, PCDHGA2, PCDHGA1, PCDHGB1, SLC25A2, TAF7 |
| WM PC1 | global measure | 10 | 50 | 133499.307 | 133541.881 | 5.00E-07 | 22 | rs12765115, rs4897813, rs9419689, rs10872824, rs9419617, rs4897759, rs9664915, rs4387283, rs4394761, rs10872825, rs7089019, rs9419618, rs9419698, rs11156567, rs10872828, rs9419702, rs7083395, rs4367873, rs9419624, rs7906770, rs9419568, rs9419569 | N/A |
| WM PC1 | global measure | 10 | 100 | 133499.307 | 133592.726 | 5.00E-07 | 35 | rs12765115, rs4897813, rs9419689, rs10872824, rs9419617, rs4897759, rs9664915, rs4387283, rs4394761, rs10872825, rs7089019, rs9419618, rs9419698, rs11156567, rs10872828, rs9419702, rs7083395, rs4367873, rs9419624, rs7906770, rs9419568, rs9419569, rs9419646, rs4243514, rs9419583, rs9419647, rs9419651, rs9419652, rs7898015, rs4307646, rs7095978, rs4243517, rs4897770, rs11156470, rs10782370 | CLVS1 |
| WM PC1 | global measure | 14 | 100 | 89530.904 | 89626.554 | 5.00E-07 | 33 | rs1033875, rs2761413, rs8006918, rs1998551, rs8017472, rs1570604, rs17714667, rs4904500, rs7144554, rs2401788, rs11621824, rs4904502, rs753361, rs4904507, rs8021839, rs10133593, rs1957841, rs4904509, rs10133171, rs4904510, rs4904512, rs927617, rs10144918, rs8015116, rs1121566, rs17125361, rs1287611, rs1018593, rs11627240, rs1475243, rs1980726, rs932731, rs11159889 | FOXN3, LOC105370615 |
| WM_09 | Superior longitudinal Fasciculus | 6 | 100 | 170585.755 | 170665.492 | 1.00E-06 | 20 | rs9348305, rs9366197, rs9348306, rs9460103, rs2180052, rs1028489, rs1033583, rs9356634, rs714869, rs2024694, rs6917485, rs9348266, rs7774272, rs9356590, rs6909300, rs9459991, rs910425, rs910424, rs11752174, rs9356648 | DLL1, FAM120B, MIR4644, LINC01624, LOC285804, LOC112267971 |

Window onset and offset are based on the reference assembly sequence GRCh37.
Any gene that overlaps with the genomic window is listed in the related gene coloumn.
